# Vitamin B_12_ alleviates Verheij syndrome-like defects via phospholipid remodeling in a *C. elegans* PUF60 spliceosomopathy model

**DOI:** 10.1101/2025.09.29.678827

**Authors:** Wenming Huang, Jonathan Kölschbach, Hormos Salimi Dafsari, Emily Baum, Anna Löhrke, Chun Kew, Ivan Dikic, Adam Antebi

**Affiliations:** Max Planck Institute for Biology of Ageing, 50931 Cologne, Germany; Cologne Excellence Cluster on Cellular Stress Responses in Aging-Associated Diseases (CECAD), University of Cologne, 50931 Cologne, Germany; Department of Pediatrics, Faculty of Medicine and University Hospital Cologne, University of Cologne, 50937 Cologne, Germany; Center for Rare Diseases, Faculty of Medicine and University Hospital Cologne, University of Cologne, 50937 Cologne, Germany; Buchmann Institute for Molecular Life Sciences, Goethe University, 60438 Frankfurt, Germany; Institute of Biochemistry II, Faculty of Medicine, Goethe University, 60590 Frankfurt, Germany; Clinical Genetics & Genomic Medicine, University Hospital of Würzburg, 97080 Würzburg, Germany

## Abstract

Verheij syndrome (VRJS) is a rare genetic disorder caused by mutations in the poly(U)-binding splicing factor 60 (PUF60), a core component of the spliceosomal complex. VRJS triggers multiple congenital anomalies, but the underlying pathomechanisms remain poorly understood. Mutation of the *Caenorhabditis elegans PUF60* ortholog, *rnp-6*, recapitulates several hallmarks of VRJS, including growth delay and smaller body size. Here, we demonstrate that developmental defects in *rnp-6* mutants are rescued by dietary K12-type *Escherichia coli* strains. Through complementary genetic screens and multi-omics analyses, we identify vitamin B_12_ (VB12) as a potent suppressor of these defects, acting via the methionine/S-adenosylmethionine/phosphatidylcholine metabolic axis. Mechanistically, *rnp-6* mutation causes aberrant splicing with methionine and phospholipid metabolism-related genes, which cumulatively impair cellular methylation potential, dysregulate phosphatidylcholine metabolism, and induce integrated stress response. We identify intron retention of the *nhr-114/HNF4* transcription factor as a primary driver of growth defects, and restoring its splicing robustly suppresses these phenotypes. VB12 supplementation bypasses the aberrant splicing, restores metabolic balance, and activates mTORC1 to rescue developmental phenotypes. Finally, we show that PUF60 deficiency induces aberrant splicing of methionine and phospholipid metabolism-related genes in a human cell line, and is associated with altered plasma methionine and phospholipid levels in VRJS patients. Our findings establish *C. elegans* as a tractable model for VRJS and uncover SAM/SAH/phospholipid dysregulation as a key mechanism underlying the spliceosomopathy, suggesting VB12 as a potential strategy to mitigate VRJS-related anomalies.

## Main

Splicing is a fundamental regulatory process in eukaryotic messenger RNA (mRNA) maturation that removes intronic regions from precursor transcripts to produce mature mRNAs encoding proteins of enhanced diversity. This essential activity is catalyzed by the spliceosome, a highly dynamic megadalton complex composed of five small nuclear ribonucleoprotein particles (snRNPs) and numerous associated splicing factors^1^. Dysregulation of splicing impairs proper cellular function and is associated with various pathological conditions, including cancer^2^ and ageing^3^. Pathogenic mutations in genes encoding core spliceosomal components disrupt splicing, leading to various genetic diseases with overlapping phenotypes^4^. *PUF60* (also known as FIR or Hfp) encodes an essential splicing factor containing two central RNA recognition motifs and a C-terminal U2AF-homology motif^5^. It operates early in spliceosome assembly by promoting the recruitment of precursor transcripts to the U2 snRNP complex and facilitating the splicing of weak 3’ acceptor sites^5,6^. Heterozygous *de novo* variants causing PUF60 deficiency result in Verheij syndrome (MIM#615583)^7,8^, a congenital disorder associated with growth retardation, short stature, and recurrent infections^8–14^. We have previously reported on a phenotypic spectrum of *PUF60*-related disorders ranging from neurodevelopmental delay to multisystem involvement^15^. However, the exact pathomechanisms of VRJS remain elusive, and no targeted therapies currently exist.

Vitamin B_12_ (also known as cobalamin) is a complex water-soluble organic compound essential for several biological functions in animals^16^. It serves as a cofactor for two metabolically important enzymes: methionine synthase (MS), which converts homocysteine to methionine within the methionine cycle, and methylmalonyl-CoA mutase, which converts L-methylmalonyl-CoA to succinyl-CoA in propionate metabolism. These two pathways are linked through homocysteine as a shared intermediate. Consequently, VB12 deficiency profoundly disrupts metabolic homeostasis and instigates various disorders in humans, including growth delay, hypotonia, anemia and cognitive impairment^16^.

In *Caenorhabditis elegans, rnp-6* (RNA-binding domain-containing protein 6) encodes the sole ortholog of human *PUF60*^17^. Depletion of *rnp-6* by RNAi-mediated knockdown causes severe developmental defects, including growth retardation, smaller body size, dysregulated immune function, and neuronal abnormalities reminiscent of VRJS in humans^18–21^. Previously, we identified a viable hypomorphic allele, *rnp-6(G281D),* that enhances abiotic stress resistance and extends lifespan on the standard B-type *Escherichia coli* OP50 diet^22,23^. In this study, we unexpectedly found that K12-type *E. coli* can fully rescue growth defects observed in *rnp-6(G281D)* mutants. Through unbiased genetic screens in *E. coli* and *C. elegans*, we discovered that *E. coli* K12 exerts its benefits through VB12-dependent methionine metabolism. VB12 restores *rnp-6* mutant growth by driving synthesis of methionine (Met), S-adenosylmethionine (SAM) and phosphatidylcholine (PC), while inhibition of enzymes required for the Met/SAM cycle abrogates such rescue. Metabolomic and lipidomic analyses reveal that *rnp-6* mutants maintained on OP50 harbor deficiencies in methylation capacity and phosphatidylcholine lipids, which are replenished by provisioning K12-type *E. coli* or VB12. Mechanistically, we deduce that aberrant splicing of *nhr-114 (*homolog of human *HNF4*), is a major proximal cause of metabolic dysregulation and developmental defects in the *rnp-6* mutant. We further demonstrate that altered Met/SAM/PC metabolism in *rnp-6(G281D)* modulates ISR and mTOR signaling. Finally, we provide evidence that PUF60 deficiency induces alternative splicing of genes related to methionine and phospholipid metabolism, and *PUF60* pathogenic mutations induce metabolic and lipidomic changes in VRJS patient plasma. Taken together, our findings implicate dysregulated S-adenosylmethionine/ S-adenosylhomocysteine (SAH) balance and phospholipid metabolism as key contributors to *rnp-6* deficiency-related growth defects, and suggest VB12 supplementation as a potential treatment to alleviate these defects.

## Results

### A K12-type *E. coli* diet alleviates growth defects in a *C. elegans* Verheij syndrome model

Previously, we identified a hypomorphic mutation in *rnp-6* that extends lifespan in *C. elegans*^22^. Further characterization showed that this mutant exhibits moderate but significant developmental defects, including smaller body size and slower growth rate when raised on the standard B-type *E. coli* OP50 diet (O) at 20°C (Figure 1a–c). These phenotypes are analogous to the growth delay and small stature seen in VRJS patients^7^. In the course of our studies, we discovered that *rnp-6(G281D)* developmental phenotypes were remarkably restored by feeding worms with the *E. coli* K12 strains HT115 (H) or BW25113 (B) (Figure 1a–c). These K12-type *E. coli* strains also suppressed other previously observed *rnp-6(G281D)* traits, including cold tolerance (Figure S1a) and extended lifespan (Figure 1d). Of note, dietary mixtures of O + B containing as little as ∼10% BW25113 sufficed to rescue growth defects (Figure S1b). Rescue did not require active bacterial metabolism, as UV-killed BW25113 also effectively restored body size (Fig. S1c). These results imply that BW25113-derived nutrients or metabolites compensate for RNP-6(G281D) dysfunction. To explore whether K12-type *E. coli* restored RNP-6 protein^20^ directly or acted downstream, we measured steady-state RNP-6 levels. However, neither HT115 nor BW25113 diets restored RNP-6 protein levels (Figure S1d–e), indicating that the rescue occurs via pathways downstream of RNP-6.

**Figure 1.**
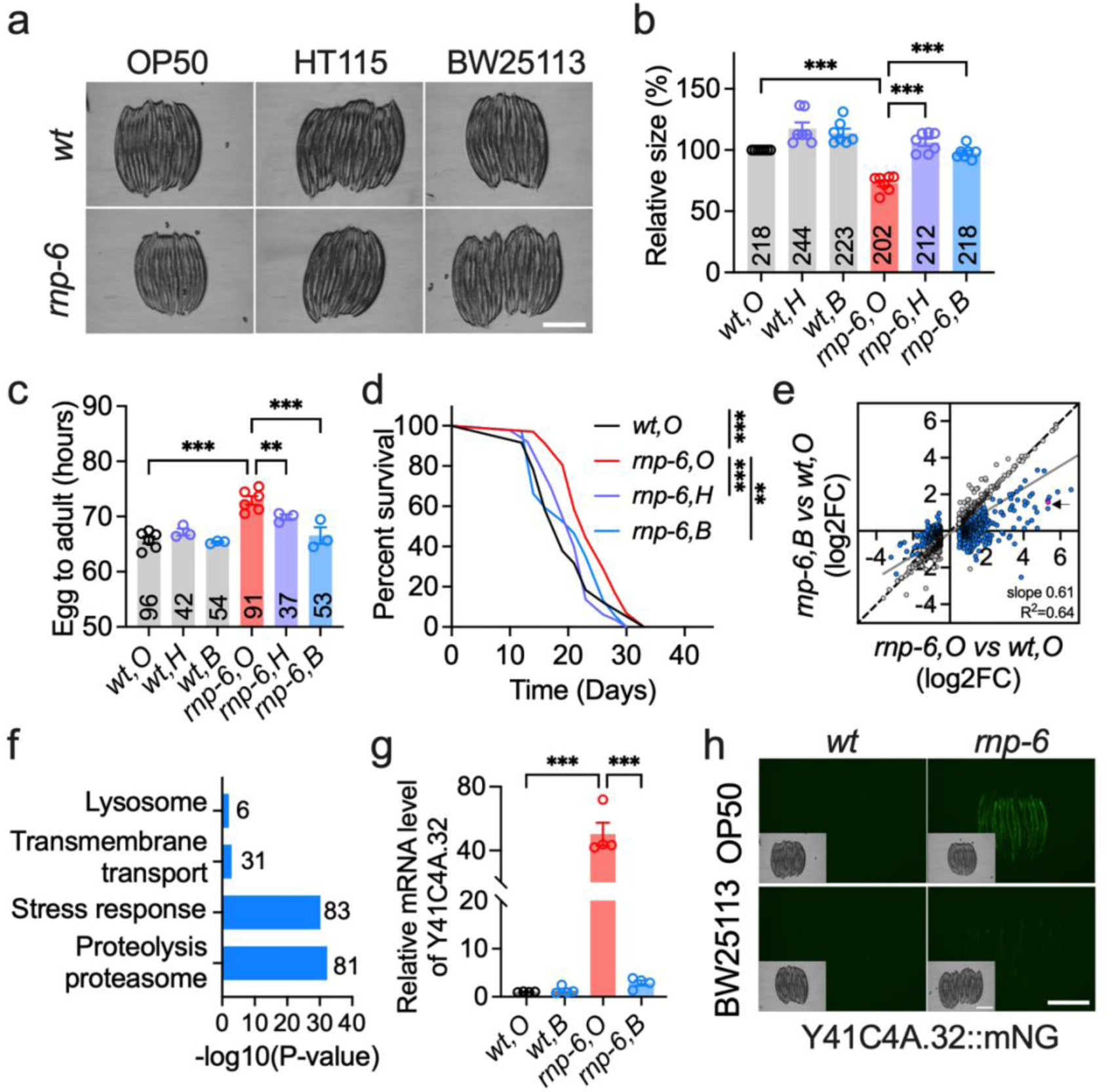
A bacterial *E. coli* K12 diet mitigates growth defects in an *rnp-6*/VRJS *C. elegans* model. **a, b**, Body size (area) of worms fed OP50 (O), HT115 (H) and BW25113 (B) (*n* = 7, scale bar: 500 μm). **c**, Assessment of K12 diet on worm growth rate (*n* = 3–6). **d**, Lifespan analysis, with survival curves illustrating one representative replicate (*n* = 3). Details of worm numbers and statistical analyses for each experimental repeat are available in Supplementary Table S16. **e,** Transcriptomic changes induced by BW25113 *E. coli*; 1541 differentially expressed genes (DEGs) regulated by *rnp-6(G281D)* relative to wild-type on OP50 (O) are shown. Blue circles indicate genes that are significantly restored by the BW25113 diet; the black arrow highlights *Y41C4A.32* expression (red circle). **f**, WormCat gene set enrichment analysis of BW25113-responsive genes. **g**, RT-qPCR quantification of *Y41C4A.32* expression (*n* = 4; F.C., fold change). **h**, Representative images of Y41C4A.32::mNeonGreen (mNG) reporter expression across the indicated diets and genotypes (n = 3, scale bar: 500 μm). Data are presented as mean ± s.e.m. unless otherwise indicated; “*n*” denotes experimental replicates, with animal counts summarized in each respective bar. Statistical analyses were conducted using the log-rank (Mantel-Cox) test for lifespan (**d**) and one-way ANOVA with Dunnett’s multiple comparisons for body size, developmental progression, and gene expression (**b**, **c**, **g**). Significance is represented as *P < 0.05, **P < 0.01, ***P < 0.001, ns: not significant. All the source data for figures are provided in supplementary tables.

To further elucidate the impact of K12-type *E. coli*, we performed RNA sequencing (RNA-seq) on wild-type and *rnp-6(G281D)* worms raised on either OP50 or BW25113 (Figure S1f). Consistent with prior work, *rnp-6(G281D)* mutants grown on OP50 exhibited profound transcriptomic changes, with 1,541 genes showing differential expression (DEseq2, adjusted p-value < 0.05, |log_2_FC| > 0.5; Table S1) and 638 splicing events in 473 genes being significantly perturbed (SAJR and IRFinder, adjusted p-value < 0.05, ΔPSI > 5%; Table S2, Figure S1g and i). About 11% of differentially spliced genes also showed altered expression (Table S3, Figure S1j), indicating distinct layers of regulation by *rnp-6*. Feeding with BW25113 significantly affected 1319 gene expression changes in *rnp-6* mutants and restored 573 genes toward wild-type levels (Figure 1e, Figure S1h, Table S4). WormCat 2.0^24^ gene set analysis of these 573 BW25113-restored genes revealed significant enrichment in stress response, proteolysis/proteasome, and transmembrane transport pathways (Figure 1f). In contrast, BW25113 diet had minimal effect on splicing profiles, with no significant restoration observed (Table S2, Figure S1k). Validation of two established *rnp-6-*dependent splicing targets, *tcer-1* and *tos-1,* confirmed this observation (Figure S1l–m). Together, these findings suggest that K12-type *E. coli* alleviates *rnp-6(G281D)* phenotypes through mechanisms downstream of, or independent from, splicing activity.

Analysis of our RNA-seq data revealed that the expression of *Y41C4A.32*, an ortholog of human COPI coat complex subunit beta 2 (*COPB2*), was among the strongest upregulated by *rnp-6(G281D)* mutation, and conversely suppressed by *E. coli* K12 (Figure 1e). This gene, previously annotated as *Y41C4A.11*, has been commonly used to assess innate immune and endoplasmic reticulum (ER) stress responses^25–28^. *rnp-6(G281D)* did not cause significant splicing changes in *Y41C4A.32* mRNA (Table S2). Quantitative RT-PCR confirmed a ∼40-fold increase in *Y41C4A.32* transcript levels on OP50 diet, which was almost completely reversed by BW25113 (Figure 1g). We thus purposed *Y41C4A.32* as a robust marker for dissecting bacteria-host interactions. To this end, we generated an endogenous C-terminal mNeonGreen fusion via CRISPR/Cas9 genome editing. Consistent with our RNA-seq and RT-qPCR results, *Y41C4A.32::mNeonGreen* expression increased by ∼10-fold in *rnp-6(G281D)* mutants compared to wild-type controls on OP50, but was substantially suppressed by BW25113 (Figure 1h). Expression of transgenic wild-type *rnp-6* fully normalized *Y41C4A.32* levels in *rnp-6(G281D)* mutants, confirming *Y41C4A.32* expression as a faithful readout of activity downstream of RNP-6 (Figure S1n–o).

### Complementary genetic screens identify vitamin B_12_ as a key alleviator of *rnp-6(G281D)* phenotypes

Utilizing the *Y41C4A.32* reporter strain, we performed a two-way genetic screen to identify K12-type *E. coli*-derived nutrients/metabolites and their corresponding host effectors (Figure 2a). We took advantage of the Keio *E. coli* mutant library, which encompasses two independent single-gene knockout mutants for 3,985 nonessential genes in the BW25113 background^29^. We posited that deletion of genes crucial for the rescue metabolite production or transport would specifically de-repress (i.e., reactivate) *Y41C4A.32* reporter expression in *rnp-6* mutants maintained on *E. coli* K12 diet. After two rounds of screening a total of 7970 Keio strains, we confirmed nine bacterial mutants that enhanced *Y41C4A.32* expression relative to the parental strain (Figure 2b). These candidate genes are involved in various biological processes, including cytoskeleton formation (*yfgA*), membrane transport (*tonB*, *btuF* and *znuB*), and purine metabolism (*purA*) (Table S5). Notably, both BtuF and TonB are core components of the *E. coli* cobalamin uptake system^30^ (Figure 2c), suggesting that VB12 or related metabolites mediate the rescue effect. Subsequent phenotypic analysis revealed that deletion of *btuF* or *tonB* in BW25113 also mimics OP50 *E. coli* effects, resulting in smaller body size of *rnp-6* mutants (Figure S2a–b). In contrast, deletion of other putative VB12 transporters (*btuB*, *btuC* and *btuD*) had negligible effects (Figure S2a–b), indicating *btuF* and *tonB* act as rate-limiting components in VB12-associated rescue.

**Figure 2.**
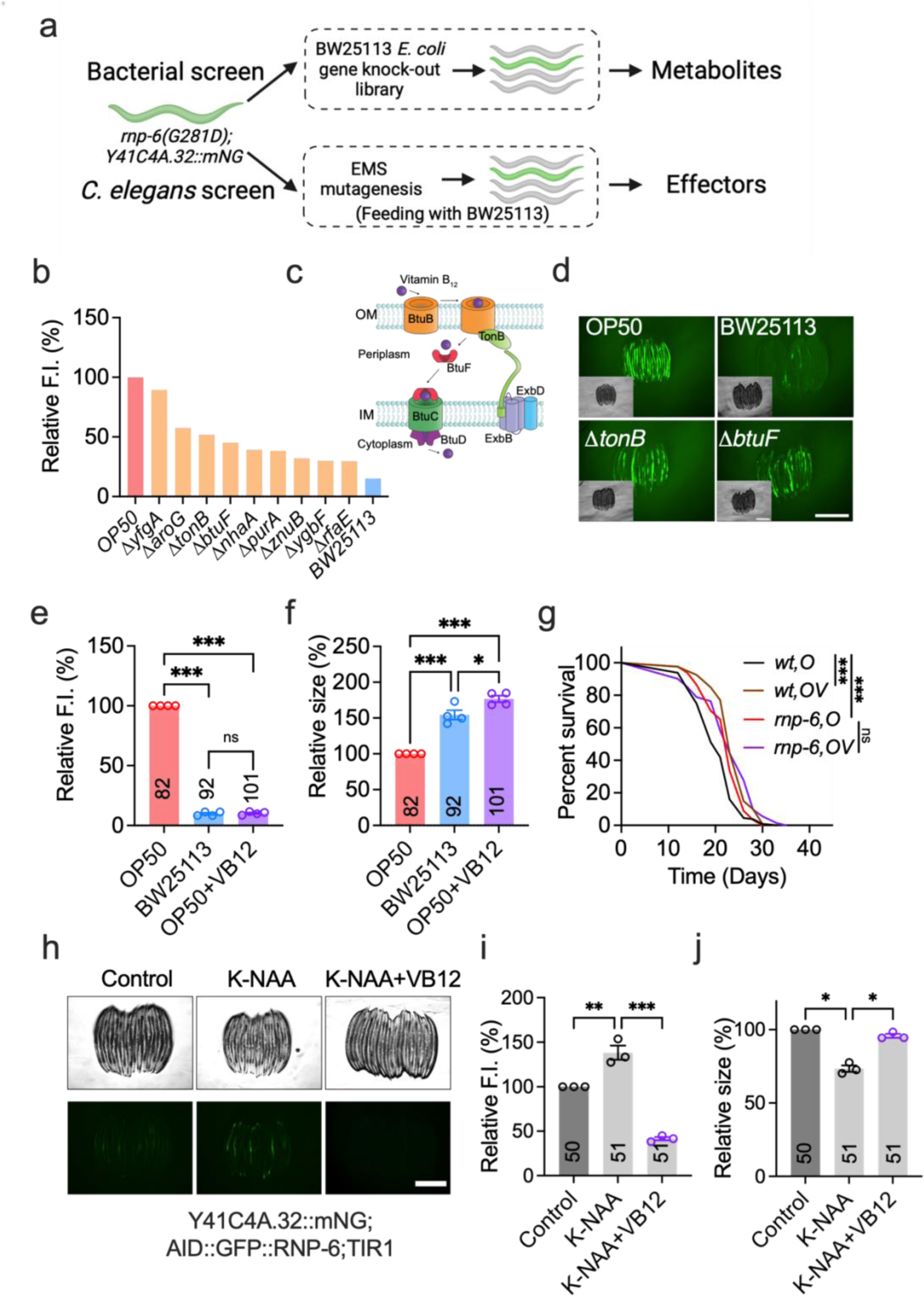
A complementary two-way genetic screen identifies vitamin B_12_ as an alleviator of *rnp-6(G281D)* mutant defects. **a**, Workflow outlining the genetic screening approach. **b**, Identification of positive hits from the Keio library *E. coli* BW25113 deletion strains. **c**, Schematic of key genes involved in VB12 uptake in *E. coli*. OM, outer membrane. IM, inner membrane. **d**, Representative images of Y41C4A.32 reporter expression in worms fed bacterial deletion mutants (*n* = 4, scale bar: 500 μm). **e–f**, Effects of VB12 (1 nM) supplementation on *rnp-6(G281D)* mutant phenotypes, shown for reporter expression (**e**) and body size (**f**) (*n* = 4). **g**, Lifespan analysis following VB12 supplementation (OV, 1 nM), with survival curves illustrating one representative replicate (*n* = 4). Details of worm numbers and statistical analyses for each experimental repeat are available in Supplementary Table S16. **h–j**, Impact of VB12 supplementation on RNP-6-depleted worms on OP50 (0.5 μM K-NAA, 1 nM VB12) (*n* = 3, scale bar: 500 μm). Statistical analyses were performed using log-rank (Mantel-Cox) survival test (**g**) and one-way ANOVA with Dunnett’s multiple comparisons (**e–f**, **i–j**). Significance is represented as *P < 0.05, **P < 0.01, ***P < 0.001. Images of **a** and **c** were generated with BioRender with permision.

*E. coli* strains cannot synthesize VB12 and largely rely on environmental scavenging^31^. Previous studies have shown that OP50 accumulates less VB12 than *E. coli* K12^32,33^, providing a plausible explanation for the strain-specific rescue of *rnp-6(G281D)*. Accordingly, we hypothesized that VB12 supplementation of OP50 would mimic the beneficial effects of a K12-type *E. coli* diet. Indeed, adding 1 nM VB12 throughout development to *rnp-6(G281D)* mutants fed on OP50 suppressed *Y41C4A.32* reporter expression and restored body size to levels comparable with BW25113-fed worms (Figure 2e–f, Figure S2c–d). VB12 rescued *rnp-6(G281D)* phenotypes even when cultured on UV-killed OP50, demonstrating a direct effect on the worm, independent of bacterial metabolism (Figure S2e–f). VB12 also alleviated other *rnp-6(G281D)* phenotypes, including slower growth rate, cold tolerance, and reduced fecundity (Figure S2g–i), confirming its role as the active metabolite mediating rescue. Surprisingly, VB12 supplementation did not reverse the *rnp-6(G281D)* longevity phenotype (Figure 2g), indicating that *E. coli* K12-derived and VB12-mediated effects on lifespan are separable. To determine the window of action, we supplemented worms with VB12 at different developmental stages. We found that VB12 treatment before the L4 larval stage fully restored normal development, matching the efficacy of continuous treatment (Figure S2j–k). Even a brief 24-hour treatment from the young adult stage onwards significantly increased body size and suppressed reporter expression (Figure S2j–k), underscoring VB12’s potent effect on *rnp-6(G281D)* development.

Given that *rnp-6(G281D)* is a specific hypomorphic allele, we investigated whether VB12’s benefits extend to broader forms of RNP-6 insufficiency that mimic the partial loss-of-function features of *PUF60* pathogenic variants in Verheij syndrome. To this end, we engineered an inducible RNP-6 loss-of-function strain using the auxin-inducible degradation (AID) system (Figure S2l)^34^. Consistent with previous data showing that *rnp-6* null mutants are embryonic lethal and complete knockdown in early life triggers larval arrest^35–37^, increasing auxin (K-NAA) doses yielded graded phenotypes and recapitulated VRJS-like defects in our system: 0.5 μM K-NAA reduced RNP-6 levels and body size (Figure S2m–n), whereas doses at 4 µM or above caused larval arrest (Figure S2n–o). Of note, partial depletion (0.5 μM K-NAA) activated the *Y41C4A.32* reporter and impaired growth (Figure 2h). Under these graded conditions of RNP-6 deficiency, VB12 supplementation restored body size and suppressed reporter induction (Figure 2h–j). These results support the notion that both *rnp-6(G281D)* and the AID-mediated partial depletion model pathogenic PUF60 haploinsufficiency, pointing to VB12 as a potential metabolic intervention.

### Vitamin B_12_ rescues *rnp-6(G281D)* mutant defects through methionine metabolism

Meanwhile, we performed EMS mutagenesis screens to identify host factors mediating rescue by K12-type *E. coli* (Figure S3a). As with the bacterial screen, we reasoned that mutations disrupting key host genes would de-repress *Y41C4A.32* expression in *rnp-6(G281D)* mutants fed BW25113. After screening ∼20,000 haploid genomes, we isolated 84 mutants that suppressed the rescue efficacy of BW25113 on reporter expression (Figure 3a). Through Hawaiian SNP mapping and whole-genome sequencing^38^, we identified causative genes for 20 mutants. Excitingly, three of these genes, *pmp-5*, *mtrr-1*, and *mthf-1,* are directly involved in the VB12-dependent methionine synthesis cycle (Figure 3b–c). *pmp-5* encodes the ortholog of human ABCD4 vitamin B_12_ transporter^39^, *mtrr-1* encodes the MTRR methionine synthase reductase^40^, and *mthf-1* encodes the MTHFR methylene-tetrahydrofolate reductase^40^. All three genes have been implicated in the “vitamin B_12_-mechanism-II” transcriptional response that senses and compensates for perturbed Met/SAM cycle activity^41^. Hence, convergent *E. coli* and *C. elegans* genetic screens strongly implicate VB12-dependent methionine metabolism as a critical modulator of *rnp-6(G281D)* phenotypes. To further validate this, we directly tested whether VB12-dependent enzymes were required for the rescue effect of BW25113 diet and VB12. Vitamin B_12_ serves as a known cofactor for two enzymes: methionine synthase (*metr-1*), which catalyzes the conversion of homocysteine to methionine, and L-methylmalonyl-CoA mutase (*mmcm-1*), which converts L-methylmalonyl-CoA to succinyl-CoA^40^. We found that the deletion of *metr-1*, but not *mmcm-1*, completely abolished the benefits of BW25113 and VB12 on *rnp-6(G281D)* mutants (Figure 3d–e). As expected, supplementation with the *metr-1* downstream product, methionine, was sufficient to bypass the requirement for *metr-1* (Figure 3f–g). Altogether, these findings suggest that *rnp-6* mutants grown on OP50 are deficient in methionine metabolism and/or downstream pathways, which can be restored by stimulating VB12-dependent methionine production. However, additional mutation of methionine biosynthesis pathway components reinstates methionine deficiency, which is rescuable by (exogenous) methionine itself.

**Figure 3.**
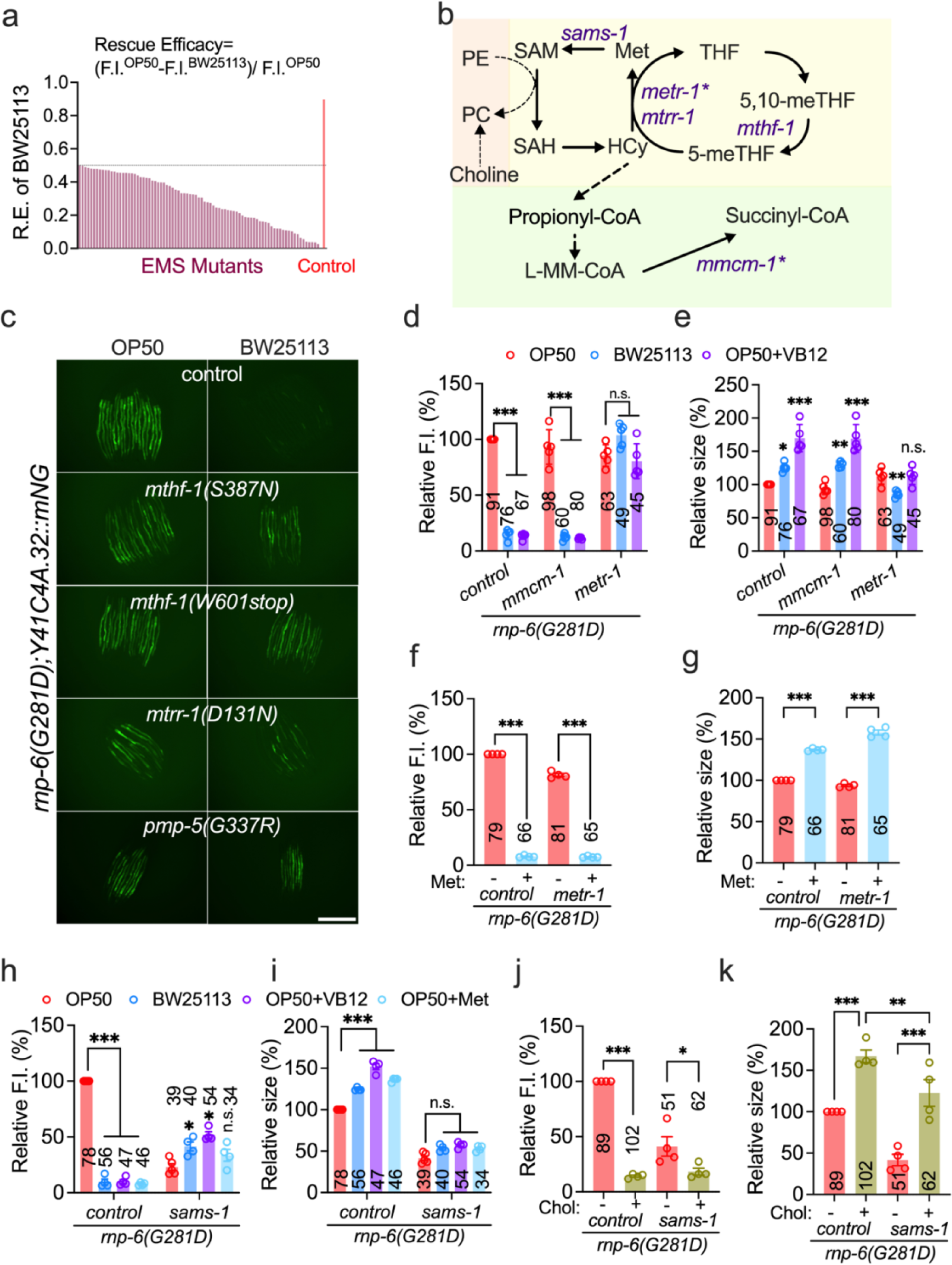
Vitamin B_12_ alleviates *rnp-6(G281D)* mutant defects through methionine metabolism pathways. **a**, Summary of hits from the *C. elegans* EMS mutagenesis screen; shown are mutants exhibiting BW25113 rescue efficacy below 50%. **b**, Schematic illustrating VB12-dependent metabolic pathways in *C. elegans*, with genes denoted by an asterisk indicating VB12 dependency. **c**, Representative images of Y41C4A.32 reporter expression in *C. elegans* mutants of EMS candidate genes (*n* = 2, scale bar: 500 μm). **d–e**, Effects of *mmcm-1* and *metr-1* deletion on reporter expression (**d**) and body size (**e**) in *rnp-6(G281D)* mutants under varying treatment conditions (*n* = 5). **f–g**, Effect of methionine supplementation (10 mM) on reporter expression (**f**) and body size (**g**) in *rnp-6;metr-1* double mutants (*n* = 4). **h–i**, Effect of *sams-1* deletion on reporter expression (**h**) and body size (**i**) in *rnp-6(G281D)* mutants under different treatment conditions (*n* = 4–5). **j–k**, Effect of choline supplementation (10 mM) on reporter expression (**j**) and body size (**k**) in *rnp-6;sams-1* double mutants (*n* = 4). Statistical significance was determined using one-way ANOVA with Dunnett’s multiple comparisons. Significance is represented as *P < 0.05, **P < 0.01, ***P < 0.001, n.s.: not significant.

We next asked how pathways downstream of methionine production interact with *rnp-6* mutant phenotypes. An important product of the methionine cycle is SAM, the principal methyl donor produced by SAM synthetase (*sams-1*)^42^. We found that deletion of *sams-1* fully abolished the rescue by BW25113, VB12 and methionine in *rnp-6(G281D)* mutants (Figure 3h–i), indicating a crucial role for SAM or SAM-dependent metabolites. Among other functions, SAM is essential for the *de novo* synthesis of PC^43,44^, a key component of cellular membranes^45^. PC can also be synthesized in a SAM-independent manner via the Kennedy pathway, which utilizes dietary choline^46^ (Figure 3b). Consistent with this framework, choline supplementation reduced *Y41C4A.32* expression, rescued body size defects, and importantly, bypassed the requirement for *sams-1* in *rnp-6(G281D)* mutants (Figure 3j–k). These findings support a model in which SAM-dependent phosphatidylcholine synthesis constitutes a major downstream mechanism by which K12-type *E. coli* and VB12 rescue *rnp-6* mutant phenotypes.

### *rnp-6(G281D)* mutation disrupts methylation potential and phosphatidylcholine metabolism

Our genetic data suggest that *rnp-6(G281D)* perturbs the methionine cycle and/or PC metabolism when grown on OP50. To examine this directly, we performed metabolomic and lipidomic analyses on *rnp-6(G281D)* and wild-type worms grown on OP50, OP50 + VB12, and BW25113 (Figure S4a). Mass spectrometry (MS)-based metabolomics identified 115 metabolites spanning amino acid, nucleotide, glucose, and polyamine metabolism (Table S6). Using MetaboAnalyst^47^, we observed that *rnp-6(G281D)* grown on OP50 significantly altered the abundance of 56 metabolites (FDR < 0.05, |log_2_FC| > 0.5) (Figure 4a-b). These metabolites were enriched in TCA cycle (succinate, citrate, malate), pyrimidine metabolism (CTP, UTP, CDP), purine metabolism (AMP, IMP, GMP), as well as the one-carbon pool by folate (SAM and SAH) (Figure S4b). Notably, the levels of SAM, SAH, and the SAM/SAH ratio were significantly decreased in *rnp-6(G281D)*, while levels of methionine, homocysteine or Met/Hcy ratio remained unchanged (Figure 4c). This pattern suggests that *rnp-6(G281D)* may impact cellular methylation potential^48^ (i.e., SAM/SAH ratio) rather than steady-state concentrations of these metabolites. Both BW25113 *E. coli* and VB12 supplementation significantly increased methylation potential (Figure 4d), confirming the central role of SAM-mediated methylation deficiency in *rnp-6* mutant phenotypes. Interestingly, several other metabolites were also rescued by BW25113 and VB12 (Figure 4b, S4c, Table S6). These included intermediates in TCA cycle and gluconeogenesis (malate, phosphoenolpyruvate) (Figure S4d–e), amino and nucleotide sugar metabolites (UDP-N-acetyl-alpha-D-glucosamine, UDP-N-acetyl-D-galactosamine) (Figure S4f–g), tryptophan metabolism components (kynurenine, tryptophan) (Figure S4h–i), and mitochondrial β-oxidation-related acylcarnitines (propionylcarnitine, butyrylcarnitine) (Figure S4j–k). Further, the oxidized-to-reduced glutathione ratio, a marker for cellular oxidative stress, was significantly elevated in *rnp-6* mutants and suppressed by BW25113 and VB12 (Figure S4l-m), indicating perturbed redox balance.

**Figure 4.**
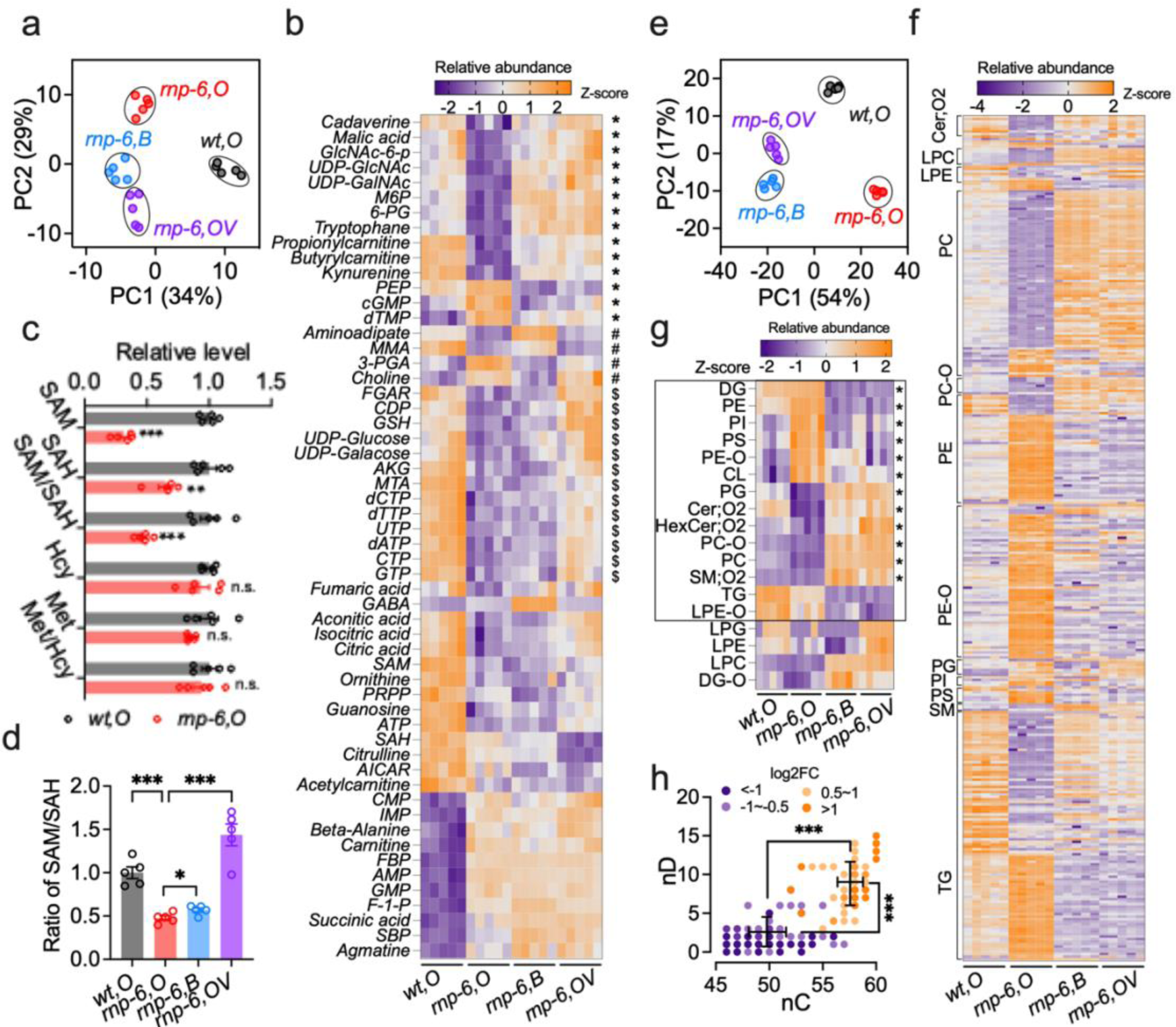
Vitamin B_12_ restores methylation potential and phosphatidylcholine metabolism in *rnp-6* mutants. **a**, Principal component analysis of metabolite profiles in wild-type and *rnp-6* mutants under different conditions (*n* = 5 per condition). **b**, Heat map of significantly altered metabolites. “*” denotes metabolites significantly restored by both BW25113 and VB12; “#” denotes metabolites significantly restored by BW25113 diet; “$” indicates metabolites significantly restored by VB12 supplementation (1 nM). The full names of all the metabolites can be found in Table S6. **c**, Relative abundance (normalized peak area) and ratio of methionine cycle-related metabolites in *rnp-6* mutants. **d**, Ratio of S-adenosylmethionine/S-adenosylhomocysteine (SAM/SAH) in *rnp-6* mutants across conditions. **e**, Principal component analysis of lipid profiles in wild-type and *rnp-6* mutants under different conditions (n = 5 per condition). **f**, Heat map illustration of significantly altered lipid species. **g**, Heat map illustration of lipid groups. The black square labels the significantly altered lipid groups in *rnp-6* mutants on OP50. “*” indicates lipid groups that are significantly rescued in by both BW25113 diet and VB12 supplementation. **h**, Distribution of the relative abundance of triacylglycerol (TG) lipid species with different fatty acyl chain lengths (nC) and numbers of double bonds (nD). TG, Triacylglycerol; Cer;O2, Dihydroceramide; LPE, Lysophosphatidylethanolamine; PG, Phosphatidylglycerol; PC, Phosphatidylcholine; LPC, Lysophosphatidylcholine; PC-O, Ether-linked Phosphatidylcholine; SM;O2, sphingomyelin; PS, Phosphatidylserine; PE-O, Ether-linked Phosphatidylethanolamine; PI, Phosphatidylinositol; PE, Phosphatidylethanolamine; DG, Diacylglycerol; LPE-O, ether-linked lysophosphatidylethanolamine; CL, Cardiolipin. DG-O, Ether-linked Diacylglycero. Statistical significance was determined by unpaired t-test (**c**, **h**) and one-way ANOVA with Dunnett’s multiple comparisons (**d**). Significance is represented as *P < 0.05, **P < 0.01, ***P < 0.001, n.s.: not significant.

MS-based lipidomics on the same sample set detected 18 lipid classes encompassing 735 species, of which the most abundant classes were phosphatidylethanolamine (PE), phosphatidylcholine (PC), and triacylglycerol (TG) (Table S7). The *rnp-6* mutation broadly altered lipid composition (Figure 4e), significantly changing the abundance of 339 lipids across 14 classes (FDR < 0.05, |log_2_FC| > 0.5) (Figure 4f-g). Notably, we found that PC were dramatically decreased, while PE were increased (Figure 4g), resulting in a significant elevation of the PE/PC ratio (Figure S5b). This pattern is consistent with impaired SAM-dependent PC synthesis in *rnp-6(G281D)* mutants. A detailed examination with individual lipids showed that TGs exhibited a biphasic response: of the 100 significantly altered TG lipids, 42% were downregulated, whereas 58% were upregulated (Figure 4f). More interestingly, the upregulated TGs contained longer and more unsaturated fatty acid chains (58 carbons, 8 double bonds on average) than downregulated TGs (50 carbons, 2 double bonds on average) (Figure 4h). Given the link between membrane phospholipid unsaturation and cold adaptation^49^, the observed lipid changes likely contribute to the enhanced cold tolerance observed in *rnp-6* mutants. BW25113 feeding and VB12 supplementation exerted similar effects on the lipidome (Figure 4e), restoring the levels of 269 and 277 lipids, respectively. 252 lipids were shared between the two treatments (Figure 4f, S5a). The relative abundance of 12 lipid classes and the PE/PC ratio were significantly rescued (Figure 4g, Figure S5b–c). These findings demonstrate that disrupted lipid metabolism is a major driver of *rnp-6* mutant defects and suggest that K12-type *E. coli* and VB12 rescue *rnp-6* defects mainly through lipid remodeling. Supporting these metabolomic findings, transcriptomic analysis revealed dysregulation of 78 lipid-metabolism genes in *rnp-6(G281D)* mutants on OP50 (Figure S5d), including peroxisomal acyl-CoA oxidases (*acox-1.2, 1.3, 1.5, 3*), fatty acid CoA synthetases (*acs-1, 2, 7, 9*), desaturases (*fat-2, 5, 6, 7*), short-chain dehydrogenases (*dhs-2, 4, 19, 23, 26, 31*), lipid-binding proteins (*lbp-5, 6, 7*), and glycerophospholipid remodeling enzymes (*ckc-1, ckb-2, eppl-1, acl-12*). Feeding BW25113 significantly restored the expression of more than half (41/78) of these genes, including *fat-2, 5, 6, 7*; *acox-1.2, 1.3, 1.5*; *acs-1,2*; *lbp-5,6*; *and ckc-1, ckb-2, acl-12* (Figure S5d).

Previous studies have shown that amino acid starvation (e.g., methionine restriction), or lipid dysregulation (e.g., PE/PC imbalance), can trigger the ISR and ER stress-related pathways^25,50–53^. Reflecting this, we detected robust phosphorylation of eukaryotic initiation factor 2α (eIF2α), an established marker of ISR activation^54^, in *rnp-6(G281D)* mutants grown on OP50 (Figure S5e–f). Consistently, the bZIP transcription factor GCN4/ATF-4, a central ISR effector^55,56^, was markedly elevated in *rnp-6(G281D)* on OP50 and effectively suppressed upon VB12 supplementation (Figure S5g–h), indicating restoration of ISR activity. In contrast, expression of *hsp-4*, a canonical ER-stress marker, and *cpl-1\**, an ER-associated degradation substrate^57,58^, was not induced by *rnp-6* mutation (Figure S4i–l). Collectively, these findings unveil a novel link from splicing dysfunction to ISR activation, rather than canonical ER stress, in *rnp-6* mutants.

### *rnp-6(G281D)* mutation causes aberrant splicing of methionine metabolism-related genes

Building on our findings of SAM and PC metabolic disruption, we sought to understand the proximal molecular mechanisms. Since RNP-6 primarily modulates pre-mRNA splicing, with potential transcriptional effects, we sought to distinguish these contributions by analyzing genetic interactions with *rbm-39*, a splicing factor associated with *rnp-6*^22,23^. We showed previously that a gain-of-function *rbm-39(S294L)* allele partially restores normal splicing and suppresses heat sensitivity of *rnp-6(G281D)* mutants^22^. Correspondingly, we found that *rbm-39(S294L)* suppressed *Y41C4A.32* expression and restored body size in *rnp-6(G281D)* mutants (Figure S6a–b). Furthermore, RNAi knockdown of splicing factors *uaf-1* and *uaf-2*, which, like *rnp-6,* are part of the U2AF complex mediating 3’ splice site recognition^5,6,59^, strongly induced *Y41C4A.32::mNG* reporter expression and reduced body size (Figure S6c). These data support a model in which *rnp-6* mutation causes stress signaling and growth defects through compromised spliceosomal functions.

To further dissect mechanisms underlying SAM and lipid dysregulation, we focused on the aberrant splicing landscape mentioned above (Figure S1j, Table S2). Among 638 significant splicing alterations, intron retention (IR) and cassette exon skipping (SE) predominated (Figure 5a). Nearly all IR events (315 out of 330 events) increased, whereas most SE events (217 out of 227 events) decreased (Figure 5b), confirming widespread disruption of canonical splicing in *rnp-6(G281D)*. Gene set enrichment analysis revealed an overrepresentation of metabolic pathway genes (Figure 5c), notably those involved in lipid (*eppl-1, haao-1, cka-1, pcyt-2.1*) and one-carbon metabolism (*metr-1, cbl-1*) (Figure S6d). We also detected an enrichment of transcription factors, including *atfs-1*, *nhr-68*, and *nhr-114*, which are involved in methionine and lipid metabolism^41,60,61^. Together, these findings implicate aberrant splicing of metabolism-related genes as a driver of *rnp-6* mutant defects.

**Figure 5.**
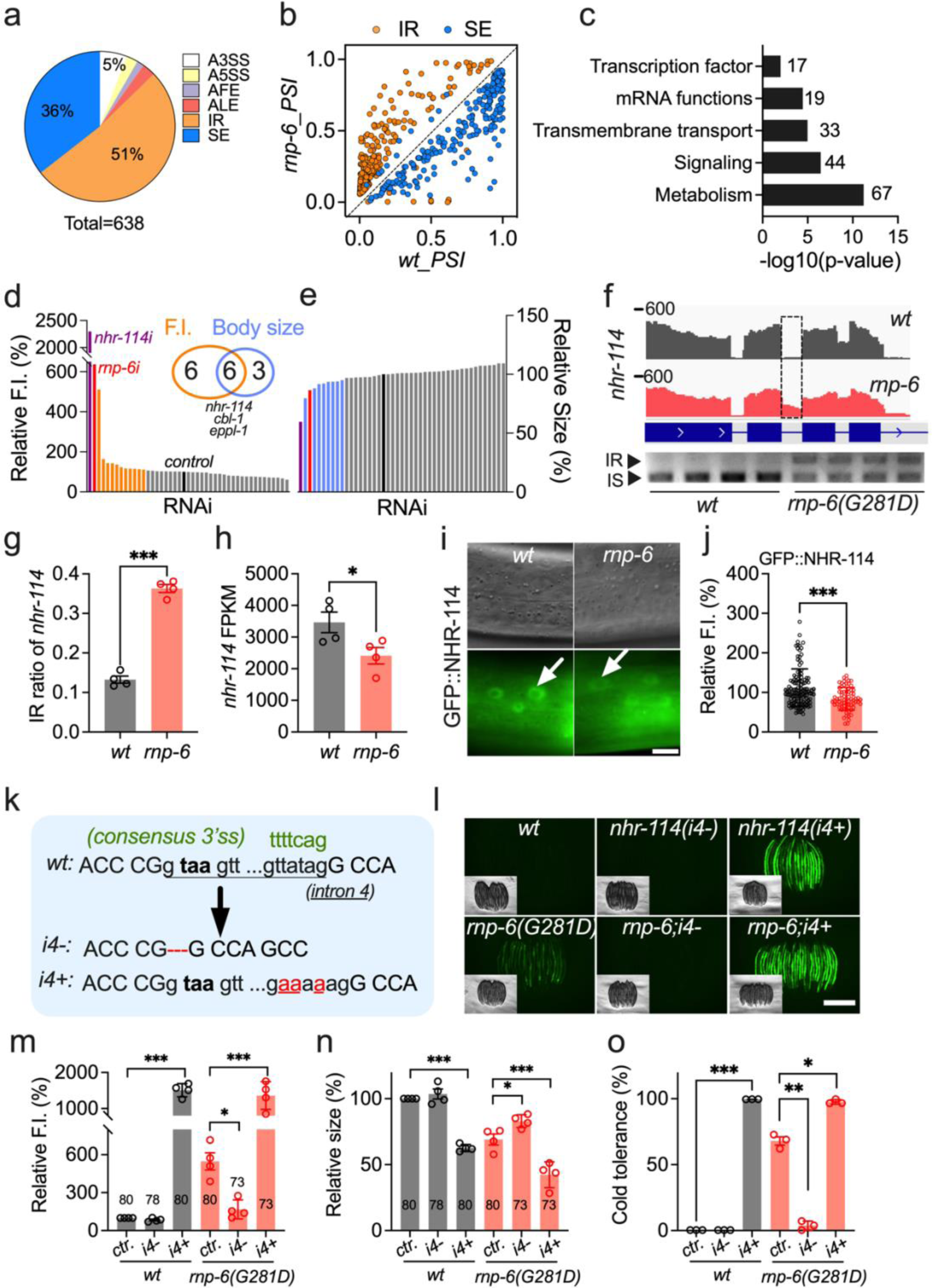
*rnp-6* mutant defects are primarily caused by aberrant splicing of transcription factor *nhr-114*. **a**, Pie chart displaying differentially spliced events in *rnp-6* mutants. A3SS, alternative 3’ splice site; A5SS, alternative 5’ splice site; AFE, alternative first exon; ALE, alternative last exon; IR, intron retention; SE, cassette exon/exon skipping. **b**, Scatter plot showing changes in cassette exon (SE) and intron retention (IR) events. **c**, Gene set enrichment analysis of differentially spliced genes using WormCat 2.0. **d–e**, Targeted RNAi screen identifying genes mimicking *rnp-6* mutant effects on Y41C4A.32 reporter expression (**d**) and body size (**e**). *luci* and *rnp-6i* were used as controls. The *mthf-1(W601stop);Y41C4A.32::mNG* strain was used to mitigate suppression effects of HT115 bacteria. **f**, Genome browser view (top) and RT-PCR analysis (bottom) of *nhr-114* intron retention (IR, intron retention; IS, intron skipping). **g**, Quantification of RT-PCR from **f** (*n* = 4). **h**, FPKM of *nhr-114* from RNA sequencing. **i–j**, Representative images and quantification of GFP::NHR-114 intensity in *rnp-6* mutant background (mean ± s.d.; scale bar: 10 μm). **k**, Schematic of CRISPR-Cas9 genome editing of *nhr-114*. The consensus sequence of intron 3 splice site is labelled in green; the red dotted line and nucleotides indicate edited residues, and bold letters in “*i4+*” represent a premature stop codon if intron 4 is retained. **l–n**, Representative images and quantification of Y41C4A.32::mNG fluorescence intensity and body size in different *nhr-114* mutants (*n* = 4, scale bar: 500 μm). **o**, Effects of *nhr-114* intron 4 editing on cold tolerance (*n* = 3). Statistical significance was determined using unpaired t-test (**g**, **h**, **j**) and one-way ANOVA with Dunnett’s multiple comparisons (**m**–**o**). Significance is represented as *P < 0.05, **P < 0.01, ***P < 0.001.

To pinpoint critical splicing targets, we filtered for metabolism-related genes and transcription factors with robust splicing changes (adjusted p-value < 0.05, ΔPSI > 0.1). This yielded 64 events, primarily involving increased intron retention or exon skipping (49/64; Table S8). Given that intron retention or exon skipping typically impairs gene function^62,63^, we conducted a targeted RNAi screen of 41 genes representing these events. In particular, we searched for candidates that phenocopied *rnp-6(G281D)*, that is, decreased body size and/or enhanced *Y41C4A.32* reporter expression in the wild-type background. In total we identified nine RNAi clones that decreased body size (>5%) and twelve that increased *Y41C4A.32* reporter expression (>10%) (Figure 5d–e; Table S9). Of those candidates, six affected both phenotypes, including *nhr-114*, *cbl-1*, and *eppl-1* (Figure 5d-e). Interestingly, these genes converge on the same metabolic network: *nhr-114* encodes an ortholog of hepatocyte nuclear factor 4 (HNF4), a key transcriptional factor regulating methionine metabolism and lipid homeostasis in *C. elegans*^41,61,64^; *cbl-1* encodes a cystathionine-beta-lyase, which converts cystathionine to homocysteine, feeding into the methionine cycle; *eppl-1* encodes an ortholog of ETNPPL ethanolamine-phosphate-phospho-lyase, which affects phosphoethanolamine metabolism.

Among these candidates, *nhr-114* showed the strongest effect on both phenotypes (Figure 5d–e, S6e–f). Deletion of *nhr-114* causes polyunsaturated fatty acid accumulation and PC depletion, both of which are reversible by VB12 or choline supplementation^61,65^. RT-PCR analyses confirmed that the *rnp-6(G281D)* mutation increased retention of *nhr-114* intron 4 by approximately 20% (Figure 5f–g). Intron 4 harbors a relatively weak 3’ splice site, and its inclusion introduces a premature stop codon (Figure 5k), likely triggering nonsense-mediated mRNA decay and reducing protein expression. Consistently, both *nhr-114* mRNA and protein levels were significantly diminished in *rnp-6(G281D)* mutants compared to wild-type controls (Figure 5h-j). Of note, *rbm-39(S294L)* gain-of-function mutation markedly suppressed intron 4 retention in *rnp-6(G281D)* mutants (Figure S6g), supporting the notion that *nhr-114* intron 4 is a direct splicing target of the RNP-6/RBM-39 complex.

To further elucidate the functional relationship between *nhr-114* and *rnp-6*, we reanalyzed publicly available RNA-seq data of the *nhr-114* null mutant^61^ and compared it with the *rnp-6(G281D)* transcriptome. As reported^61,64^, *nhr-114* deletion caused widespread transcriptional changes, altering the expression of 4,235 protein-coding genes (DEseq2, adjusted p-value < 0.05, |log_2_FC| > 0.5) (Table S10). 483 of these genes were also significantly altered in *rnp-6(G281D)* (overlap significance, p < 5e-12) (Figure S6h–i; Table S10). In particular, genes related to lipid metabolism were significantly enriched (Figure S6j), including peroxisomal acyl-CoA oxidases (*acox-1.2, 1.3*), fatty acid desaturases (*fat-2, 5, 6, 7*), lipid binding proteins (*lbp-5, 6, 7*), and phospholipid-remodeling genes (*ckb-2, acl-12*) (Table S10). Using a less stringent cutoff (adjusted p-value < 0.05, |log_2_FC| > 0.1), we also observed overlap of one-carbon cycle genes (*ahcy-1, dao-3*) (Table S10). Less than 4% (18/483) of the overlapped DEGs displayed aberrant splicing in *rnp-6* mutants (overlap significance, p < 0.102) (Table S10), supporting separable splicing and transcriptional contributions. Strikingly, ∼60% (288/483) of the shared genes were rescued by VB12 supplementation in the *nhr-114* mutant and by BW25113 feeding in the *rnp-6* mutant (Figure S6k, Table S10), suggesting common downstream pathways. These findings strongly implicate intron retention-induced *nhr-114* loss-of-function as a key mediator of *rnp-6(G281D)* transcriptomic changes and suggest that splicing and transcriptional programs converge on the Met/SAM/PC axis to cause the observed phenotypes.

To directly test this hypothesis, we generated two *nhr-114* alleles manipulating intron 4 retention: *nhr-114(i4+)*, with three thymine-to-adenine substitutions at the 3’ splice site to enhance intron retention, and *nhr-114(i4–)*, with complete intron 4 deletion (Figure 5k). RT-PCR verified the intended splicing alterations (Figure S6l). As predicted, *nhr-114(i4+)* phenocopied *rnp-6(G281D)*, reducing body size, robustly activating the *Y41C4A.32* reporter, and increasing cold tolerance in the wild-type background, while not further elevating *Y41C4A.32* expression in *rnp-6(G281D)* mutants (Figure 5m–o). Conversely, the *nhr-114(i4–)* allele alone produced no overt phenotype in wild-type animals (Figure 5m–o), but significantly suppressed multiple *rnp-6(G281D)* phenotypes, including *Y41C4A.32* reporter expression, cold tolerance, and body size defects (Figure 5m–o). Taken together, these results provide compelling evidence that intron 4 retention of *nhr-114* is a major driver of *rnp-6* mutant pathology.

Lastly, we explored the physiological regulation of *nhr-114* splicing. Dietary restriction (DR) decreases SAM levels in flies and mice^66–68^, and DR-induced longevity is mediated by *sams-1* in *C. elegans*^69,70^. Further, nutrient-sensing pathways have been shown to regulate splicing and metabolism^71^. We hypothesized that nutrient availability modulates SAM metabolism partially through *nhr-114* splicing. We therefore investigated *C. elegans* adult reproductive diapause (ARD), a state induced by long-term fasting and associated with downregulation of RNA processing complexes^72,73^. Under ARD conditions *nhr-114* intron 4 retention was significantly increased by ∼10% relative to fed controls (Figure S6n), accompanied by reduced levels of *nhr-114* mRNA (Figure S6o). These findings suggest that nutrient status modulates *nhr-114* splicing and the associated SAM/phospholipid metabolic network.

### Vitamin B_12_ activates mTOR signaling to rescue *rnp-6(G281D)* mutant defects

mTOR signaling orchestrates key processes of development, growth, and metabolism^74^. Previously, we showed that *rnp-6(G281D)* mutation inhibits mTORC1 activity when worms are fed OP50^22^. Here, we asked whether VB12 supplementation could restore mTORC1 function in these mutants. To this end, we (i) measured phosphorylation of AMPK, a cellular energy sensor inversely related to mTORC1 activity, and (ii) monitored nuclear localization of HLH-30, the *C. elegans* ortholog of TFEB, whose nuclear translocation is suppressed by active mTORC1 signaling^75–77^. Consistent with mTORC1 inhibition, *rnp-6(G281D)* mutants on OP50 exhibited elevated AMPK phosphorylation (Figure 6a–b) and increased HLH-30 nuclear localization (Figure 6c–d). Remarkably, VB12 supplementation reversed both phenotypes, indicating a reactivation of mTORC1 signaling.

**Figure 6.**
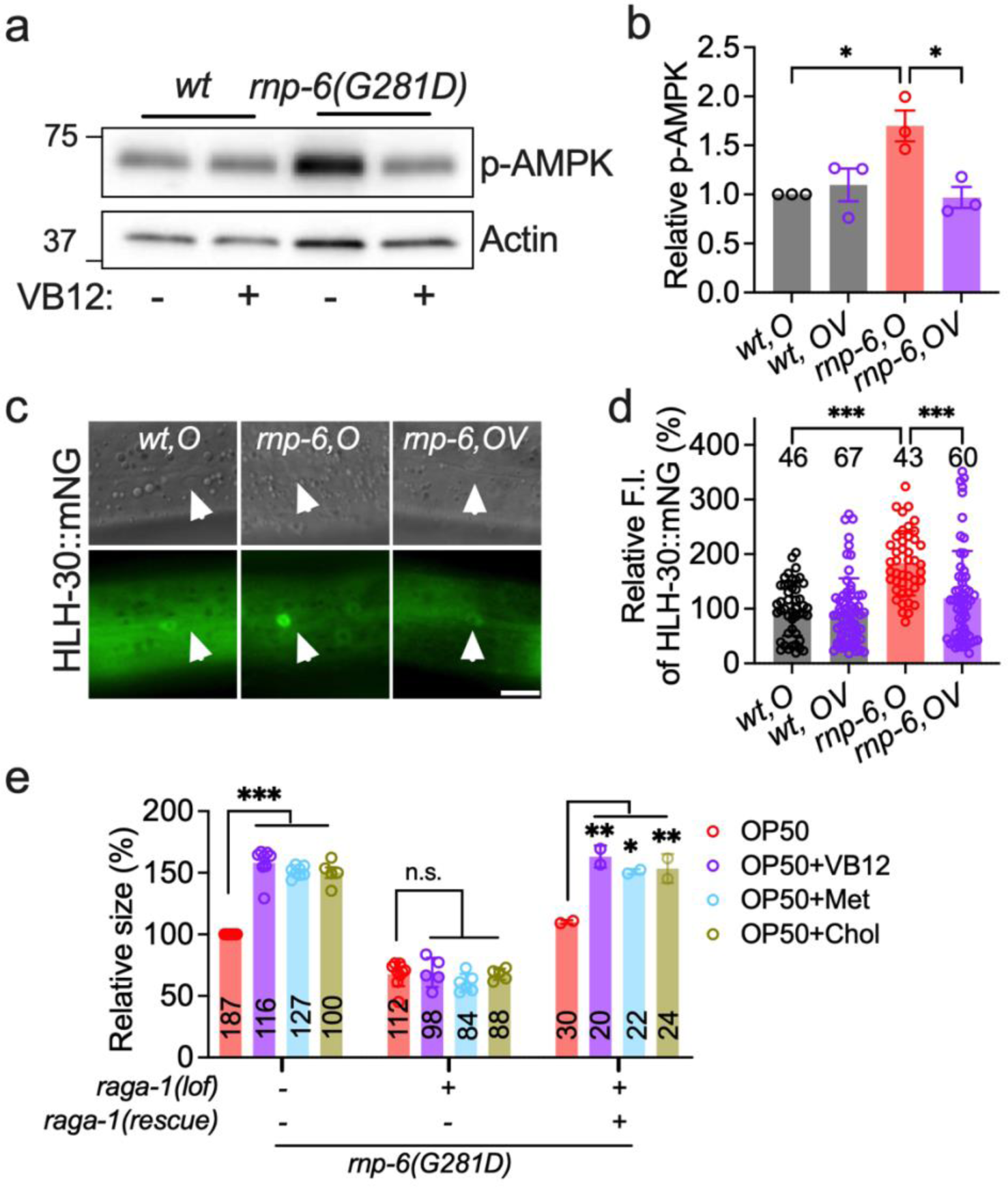
Vitamin B_12_ restored mTOR signaling in *rnp-6(G281D)* mutant. **a**, Western blotting of AMPK phosphorylation in *rnp-6* mutants. **b**, Quantification of Western blotting experiments in a. n=3. **c**, Representative images of HLH-30::mNG nuclear localization in *rnp-6* mutants (hypodermis). Scale bar, 20 mm. **d**, Quantification of nuclear HLH-30::mNG fluorescent intensity. The results from three biological replicates were merged. **e**, The effects of *raga-1* deletion and *raga-1* single-copy insertion on worm body size in *rnp-6* mutants. n=2-7. One-way ANOVA with Dunnett’s multiple comparison test were used for statistical analysis. *P < 0.05, **P < 0.01, ***P < 0.001, n.s., not significant.

Next, we wondered whether mTORC1 activity is required for rescue of *rnp-6(G281D)* growth defects by VB12/Met/SAM/PC. Loss-of-function mutation of *raga-1*, a Rag GTPase essential for mTORC1 activation, further reduced the body size of *rnp-6* mutants and completely abolished the beneficial effects of K12-type *E. coli*, as well as VB12, methionine, and choline supplementation on body size in *rnp-6(G281D)* mutants (Figure 6e). A *raga-1* single-copy insertion^77^ rescued body size in the *raga-1(lof);rnp-6* double mutant on OP50 and fully reinstated the rescue of body size upon VB12, methionine or choline supplementation (Figure 6e). Altogether, these findings demonstrate that intact mTORC1 signaling is critical for mediating the rescue effect of VB12 and related metabolic interventions in *rnp-6(G281D)* mutants.

### PUF60 deficiency impairs 1CC and phospholipid metabolism in human cells

Since RNP-6 deficiency induces aberrant splicing of 1CC and phospholipid metabolism genes in *C. elegans*, we wondered whether PUF60 deficiency has a similar effect in human cells. We mined public datasets in which the human lung adenocarcinoma cell line PC9 was subjected to PUF60 siRNA^78^. Based on our findings in worms, we reasoned that splicing defects would remain unchanged even when cells were cultured in the presence of VB12 and methionine-rich medium. We re-analyzed the data using the SAJR and IRfinder pipelines and combined the results with those generated with rMATS in the original paper. In total, we detected 2599 splicing changes corresponding to 1730 genes (adjusted p-value < 0.05, Δpsi > 0.1) (Figure S7a, Table S11). KEGG analysis of the aberrant spliced genes revealed the significant enrichment of metabolic pathways (Figure 7a, Table S12). These include genes related to propanoate metabolism (*e.g.,* ethylmalonyl-CoA decarboxylase 1 *ECHDC1/C32E8.9*, methylmalonyl-CoA epimerase *MCEE/mce-1*), choline metabolism (*e.g.,* choline kinase alpha *CHKA/cka-2*), glycerophospholipid metabolism (diacylglycerol kinase *DGKQ/dgk-1*, phosphocholine cytidylyltransferase *PCYT1A/pcyt-1*, phosphatidate phosphatase *LPIN2/lpin-1*), cysteine and methionine metabolism (methionine synthase *MTR/metr-1*, kynurenine aminotransferase 1 *KYAT1/nkat-1*) and VB12 transport (*ABCD4/pmp-5*) (Figure 7b). These results together imply that PUF60 deficiency might dysregulate 1CC and phospholipid metabolism through altered splicing.

**Figure 7.**
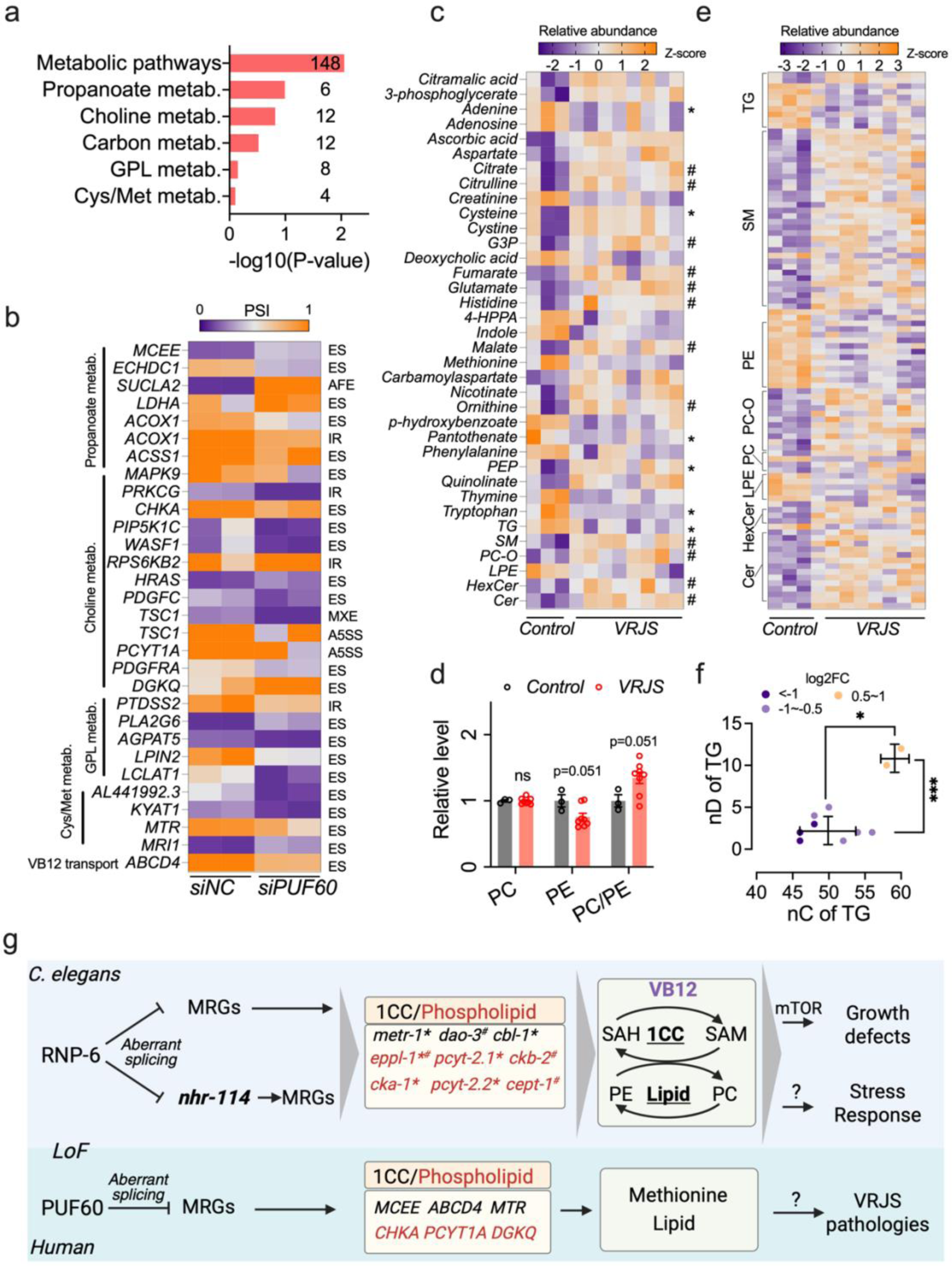
PUF60 deficiency dysregulates 1CC and phospholipid metabolism. **a**, KEGG pathway analysis of PUF60-dependent splicing events in PC9 lung cancer cell line. GPL, glycerophospholipid. **b**, Heatmap of aberrant spliced genes related to methionine and phospholipid metabolism. **c**, Heatmap illustration of significantly altered metabolites and lipid groups. “*” and “#” indicate the metabolite/lipids that show the same and opposite direction of changes, respectively, between VRJS plasma samples and worm samples. d, Quantification of the relative level of PE and PC in VRJS plasma. **e**, Heatmap illustration of altered lipids in VRJS plasma. **f**, Distribution of the relative abundance of triacylglycerol (TG) lipid species with different fatty acyl chain lengths (nC) and numbers of double bonds (nD). **g**, Model depicting the pathomechanisms underlying RNP-6/PUF60 deficiency. RNP-6 loss-of-function (LOF) causes aberrant splicing of metabolism-related genes (MRGs) as well as metabolism-related transcriptional factors such as *nhr-114,* which in turn alter the expression of MRGs. Altered MRGs collectively cause an imbalance of SAM/SAH and PE/PC metabolic axis, which triggers stress responses (including ISR and mitochondrial stress) and delays growth through inhibiting mTOR activity. VB12 supplementation restores SAM/SAH balance and PE/PC metabolism, suppressing stress response and rescuing normal development. “*” indicates the splicing targets of *rnp-6*; “#” indicates the transcriptional targets of *nhr-114*. In humans, PUF60 loss-of-function causes aberrant splicing of 1CC and phospholipid metabolism-related genes and is associated with altered levels of methionine and lipid levels in patient plasma. Dysregulated metabolism might drive some pathologies in VRJS patients.

Last, we sought to understand if the *PUF60* pathogenic mutation also affects metabolism in VRJS patients. To this end, we collected blood samples from 8 Verheij syndrome patients (6 males and 2 females) and 3 age-matched non-*PUF60* mutation controls (1 male and 2 females) and performed metabolomic and lipidomic analyses with plasma (Figure S8b). In total 103 metabolites and 653 lipid species belonging to 15 lipid groups were annotated (Table S13-14). Although variable, the VRJS samples clustered differently from controls in PCA plots (Figure S7c). Out of the 103 metabolites, 30 were significantly altered (p<0.05, log2FC>0.2) (Figure 7c). Strikingly, tryptophan and methionine were among the most significantly changed metabolites, decreased by ∼90% and ∼85% in VRJS, respectively (Figure S7d-e). A handful of metabolites involved in TCA cycle, such as citrate, malate and fumarate, were upregulated in VRJS patients (Figure S7f-h). We were unable to quantify SAM or SAH as they were not detected in our plasma samples, due to low abundance. Levels of TG and LPE were significantly decreased, while the abundance of PC-O, hexose ceramide, ceramide and sphingomyelin lipid were significantly increased (P<0.05) (Figure 7c). Although the levels of PC remained unchanged, the levels of PE and the ratio of PC/PE were slightly increased (Figure 7d). When we checked individual lipids, 94 were significantly changed (p<0.05, log2FC>0.2) (Figure 7e). Consistent with the group analysis, most of the significantly altered PE and LPE were downregulated, while PC-O, sphingomyelin, ceramide and hexose ceramide were upregulated (Figure 7e). Interestingly, we found that, similarly to *C. elegans,* the significantly upregulated TG lipids contained longer and more unsaturated fatty acid chains (59 carbons, 11 double bonds on average) than downregulated TGs (48 carbons, 2.5 double bonds on average) (Figure 7f). Taken together, our results from human cell lines and patient plasma provide evidence that PUF60 deficiency might dysregulate sulfur amino acid and phospholipid metabolism.

## Discussion

Verheij syndrome remains an exceptionally rare and poorly understood genetic disorder with no targeted treatments to date. Albeit relatively simple, *C. elegans rnp-6* hypomorphic mutants recapitulate multiple features of VRJS such as smaller body size, growth delay, immune alterations, and neurological defects^7,9,10,12,14,15,79^, highlighting the nematode as a powerful system for dissecting the molecular and cellular mechanisms of PUF60 insufficiency-associated pathologies. Leveraging the *C. elegans rnp-6* mutant as an *in vivo* disease model, we found that VRJS-associated growth defects arise primarily from SAM/SAH and PE/PC imbalance. The metabolic impairments are cumulatively driven by aberrant splicing of one-carbon and phospholipid metabolism-related genes (such as *metr-1*, *cbl-1*, *eppl-1*) and *nhr-114*, a pivotal transcription factor of one-carbon and phospholipid metabolism. Reconstitution of SAM/SAH and PE/PC balance through supplementation of VB12, methionine, choline, or VB12-enriched bacteria can robustly rescue development (Figure 7g). These interventions mechanistically converge on reactivating mTORC1 signaling, a key driver of development, establishing a direct link between splicing factor dysfunction, metabolic rewiring, and nutrient-sensing signaling networks. To our knowledge, this is the first study to integrate the aforementioned metabolic dysregulation with spliceosomopathies, and to identify vitamin B_12_ as a candidate therapeutic for VRJS-like pathologies.

*rnp-6* mutation disrupts expression and splicing of diverse metabolism-related genes (Table S1 and S2). By combining targeted RNAi screening, genetic epistasis analysis, and transcriptomic profiling, we identify aberrant intron retention of *nhr-114* as a critical node linking RNP-6 perturbation to downstream transcriptional and metabolic imbalances, thereby driving defects in *rnp-6* mutants. As *nhr-114*/*HNF4* homologs govern pivotal aspects of methionine and lipid homeostasis across species^41,61,64,80,81^, our findings offer a mechanistic bridge connecting spliceosomal dysfunction with metabolic dysregulation. Moreover, the novel observation that *nhr-114* splicing is dynamically regulated by both genetic perturbation and physiological states (such as fasting-induced diapause) further broadens the connection between nutrient-responsive control of RNA processing and cellular metabolism.

VB12 is an essential micronutrient critical for human health, particularly in preventing anemia and neurological dysfunction^16^. Clinically, VB12 supplements address malabsorption disorders such as pernicious anemia and gastrointestinal disorders^82^. Beyond these classic roles, emerging studies demonstrate that VB12 enhances somatic cell reprogramming and tissue repair in mammalian models^83^. In *C. elegans*, VB12 deficiency induces severe phenotypes, including cognitive impairment, growth retardation, infertility, and shortened lifespan^84,85^. Supplementation of VB12 mitigates amyloid-beta and dithiothreitol-induced toxicity in *C. elegans*^86,87^, highlighting its essential role in metabolic homeostasis. Our work extends these insights by demonstrating that VB12 supplementation rescues RNP-6 deficiency-associated spliceosomopathy, confirming the critical function of VB12 under both physiological and disease conditions.

Beyond the central role of SAM/SAH balance and phospholipid metabolism, our data also implicate additional metabolic pathways and organelles as contributors to VRJS-like defect manifestations. For example, metabolites related to the tricarboxylic acid (TCA) cycle (e.g., malate, phosphoenolpyruvate) and mitochondrial β-oxidation (propionylcarnitine and butyrylcarnitine) are reduced in *rnp-6* mutants, implying mitochondrial dysfunction. Supporting this, the mitochondrial stress reporter *hsp-6* is strongly induced in *rnp-6(G281D)* mutants and fully suppressed by the K12-type *E. coli* strain BW25113 (Figure S8a–b). This is also consistent with previous reports that link VB12 supplementation to improved mitochondrial function^88,89^. Moreover, crucial metabolites in the *de novo* NAD^+^ synthesis pathway, such as kynurenine and tryptophan, are reduced in *rnp-6* mutants and restored by BW25113 *E. coli* or VB12, hinting at dysregulated NAD^+^ metabolism in *rnp-6(G281D)* pathology. Additionally, metabolites associated with the glucosamine pathway appear dysregulated. A unifying notion is that *rnp-6* mutation induces a catabolic state of lower methylation capacity, mitochondrial function, and glucose metabolism, while VB12 reverses these features and re-establishes a more anabolic state, consistent with the activation of mTOR signaling.

Of particular interest is the induction of the integrated stress response in *rnp-6* mutants. The ISR is a conserved pathway for eukaryotic cells that limits protein synthesis and secretion to cope with diverse stresses^54^. The impact of the splicing machinery on ISR has been little explored, though a recent investigation shows that aberrant splicing of protein translation genes induces the ISR and inflammation-related gene expression in acute myeloid leukemia patients^90^. Our results provide direct evidence that splicing factor dysfunction triggers the ISR in *C. elegans*, in this case through phospholipid remodeling. Further research should help elucidate the detailed mechanisms.

Our study demonstrates that VB12 supplementation activates mTORC1 signaling to rescue growth and metabolic defects in the *rnp-6* mutant, revealing a novel mechanistic link between micronutrient status and spliceosomal dysfunction. Previous work showed that methylcobalamin, a VB12 analog, activates mTOR via Akt to promote neuronal outgrowth, supporting VB12’s capacity to stimulate mTOR in a cellular context^91^. In contrast, recent studies report VB12 inhibits mTOR phosphorylation to induce autophagy in disease models^92^. Our data uniquely position VB12-induced mTORC1 activation as an anabolic rescue mechanism critical for correcting spliceosomal metabolic defects. Although mTOR activation typically limits longevity in *C. elegans*, the lack of lifespan limitation by VB12 may reflect tissue-specific effects of mTORC1, which restricts lifespan in neurons but promotes growth in somatic tissues^23–25^. We speculate that VB12 preferentially activates mTORC1 in non-neuronal tissues to support development without affecting longevity. Dissecting the tissue-specific roles of RNP-6/NHR-114 signaling, SAM/SAH–PC/PE metabolism, and mTOR within this metabolic network will be important to elucidate. While K12-type *E. coli* (BW25113 and HT115) and VB12 exert largely overlapping effects in *rnp-6(G281D)* mutants, the bacterial diet may also regulate physiology through VB12-independent mechanisms. For instance, K12-type *E. coli* suppresses *rnp-6(G281D)*-associated longevity (Figure 1d), whereas VB12 treatment shows little effect (Figure 2g), suggesting bacterial metabolites or lipids beyond VB12 contribute to lifespan regulation^93^. Together, our findings highlight the pleiotropic and context-dependent modulation of mTOR by VB12 and advance current paradigms by identifying mTORC1 as a promising therapeutic target in spliceosomal diseases.

Our analysis of splicing changes in the PC9 cell line suggests that RNP-6/PUF60 might play conserved roles in regulating methionine and phospholipid metabolism, as we observed enrichment of aberrantly spliced genes in choline, propanoate, and methionine/cysteine metabolism. Though we did not find (significant) splicing changes in *HNF4* (functional homolog of *nhr-114),* we did see splicing changes in *NR1H3* (Table S11) a related nuclear receptor that also regulates lipid metabolism^94^, suggesting that RNP-6 and PUF60 may affect similar and different splicing targets, dependent on cell type. It is also interesting to note that mTOR signaling pathway is significantly enriched in the PUF60-dependent splicing genes, including NPRL2, NPRL3, RPS6KB2, RPS6KA3 and TSC1 (Table S12). The aberrant splicing with mTOR complex core components might contribute to mTORC1 inhibition under PUF60 deficiency conditions^22^. By performing metabolic and lipidomic assays in VRJS plasma, we provide evidence that the PUF60 pathogenic mutation might dysregulate methionine/cysteine and phospholipid metabolism. Of note, some metabolites (such as cysteine, tryptophan, PEP, pantothenate, adenine and long chain TGs), were changed in the same direction in *rnp-6* mutants and VRJS patients, while other metabolites (such as citrate, malate and fumarate) and lipids (such as PE, hexose ceramide, sphingomyelin lipid and PC/PE ratio) were changed in the opposite direction. Despite these discrepancies, these findings suggest that similar pathways are dysregulated, while the directionality could reflect different metabolic/lipidomic profiles of plasma versus tissues^95,96^. Moreover, as the plasma samples were obtained from subjects of different ages and genders, this may also partially contribute to the observed changes. It will be of great importance to validate these results in plasma and patient-derived cells (such as fibroblasts) with a larger cohort (age and gender matched) and test the effect of VB12 supplementation.

Splicing is a highly coordinated process and involves hundreds of splicing factors^1^. RNP-6/PUF60 functionally and physically interacts with many other splicing factors that act at various stages of spliceosome assembly^97^. Mutations in some of these splicing factors, including UAF-1/U2AF2^98,99^, PRP-19/PRPF19^99^, PRP-8/PRPF8^100^, RSP-7/SRSF11^101^ and RBM-5/RBM10^102^, also trigger VRJS-like neurodevelopmental defects in humans, suggesting potential convergent pathomechanisms. Given the neurological manifestations in these spliceosomopathies and the neuronal relevance of *rnp-6*^21,22^, assessing VB12’s impact in this context represents critical future work. Most importantly, it will be of great interest to investigate if SAM/SAH or phospholipid metabolism are similarly dysregulated in these mutants and test whether VB12 can alleviate associated growth and developmental defects.

In summary, our results from a VRJS spliceosomopathy model in *C. elegans*, a human cell line and patient samples illuminate an unappreciated metabolic bottleneck underlying RNP-6/PUF60 deficiency and warrant further investigation of VB12 supplementation in treating VRJS syndromes.

## Methods

### *C. elegans* strains and maintenance

Worms were maintained at 20°C following standard procedures^103^. Detailed information for strains used in this study are found in Supplementary Table S15. All mutant strains obtained from CGC and Sunybiotech (www.sunybiotech.com) were outcrossed with our N2 at least twice. Detailed information regarding CRISPR gene editing can be provided upon request. For all experiments, worm synchronization was done by egg laying.

### Compound supplementation assay

For supplementation of methionine (L-methionine, Sigma-Aldrich: M9625) and choline (choline chloride, Thermo Fisher: A15828.22), methionine/choline was added to NGM medium before pouring to a final concentration of 10 mM. OP50 was seeded on these plates two days before worms lay eggs. For vitamin B_12_ supplementation, VB12 (Cyanocobalamin, Sigma-Aldrich: V6629) was mixed with OP50 bacteria and seeded on NGM plates. Worms were cultured on these plates from the egg stage onwards, unless noted otherwise. For the OP50-BW25113 mixture assay, OP50 and BW25113 were cultured to log phase, washed with M9 and concentrated to the same OD600 value. Then OP50 and BW25113 was mixed at different ratios and seeded on peptone-free NGM plates to avoid bacterial growth. Worm images were taken at the young adult stage. For the tunicamycin response assay, tunicamycin (Sigma-Aldrich: T7765) stock solution was added on top of seeded NGM plates (final concentration 2 ug/ml) with synchronized young adult stage worms. After overnight treatment (∼16 h), worms were anesthetized with sodium azide (50 mM) and imaged.

### Worm imaging

Analysis of worm reporters Y41C4A.32::mNeonGreen and worm size were performed on a Leica stereo microscope (Leica M165 FC, LAS X) with Leica DFC3000G CCD camera. Analysis of mTOR reporter HLH-30::mNeonGreen was performed on a Zeiss Axioplan2 microscope (Axio Vision SE64, Rel.4.9.1) with a Zeiss AxioCam 506 CCD camera. Fiji software (Version 2.0.0/1.52p)^104^ was used for quantifying fluorescent intensity and worm area. For HLH-30::mNeonGreen images, the nuclei of hypodermal cells were selected for quantification. To reduce bias, worms were randomly picked under a dissection microscope and imaged. At least 20 worms per genotype were picked for imaging and all the experiments were independently carried out at least three times unless otherwise indicated.

### Developmental rate assay

Worms were synchronized by short-term (1h) egg lay on the indicated plates. After 48 hours of growth, ∼15 worms from each condition were randomly singled to 3 cm NGM plates. After 12 hours, each worm was checked every hour until it laid the first egg. Experiments were repeated at least three times.

### Brood size assay

Worms were synchronized by short term (1h) egg lay on the indicated plates. After 48 hours of growth, ∼15 L4-stage worms of each condition were singled to 3 cm NGM plates. Worms were transferred every day until egg laying stopped. The total number of hatched larval worms was counted. Experiments were repeated at least three times.

### Cold tolerance assay

Worms were synchronized and grown on the indicated plates. When the worms reached the young adult stage, they were transferred to a 2°C incubator for 24 hours. Worms were recovered at room temperature for 4 hours and the number of alive and dead worms was scored. Cold survival ratio was measured as the ratio of the number of live worms to the number of total worms. At least 60 worms from each genotype were used in the assay for each replicate. Experiments were repeated at least three times.

### Lifespan assay

All lifespans were performed at 20°C with the indicated diet or treatment condition for the whole life. Worms were allowed to grow to the young adult stage on standard NGM plates. ∼150 young adults were transferred to NGM plates supplemented with 10 µM of FUdR. Survival was monitored every other day. Worms that did not respond to gentle touch by a worm pick were scored as dead and were removed from the plates. Animals that crawled off the plate or had ruptured vulva phenotypes were censored. All lifespan experiments were blinded and performed at least three times. Graphpad Prism (9.0.0) was used to plot survival curves. Survival curves were compared, and p-values were calculated using the log-rank (Mantel-Cox) analysis method. Complete lifespan data are found in Supplementary Table S16.

### RNA interference

RNAi experiments were performed as previously described^23^. *E. coli* HT115 (high VB12) and *E. coli OP50(xu363)* (low VB12) bacterial strains were used in this study. HT115 bacteria came from the Vidal or Ahringer library. *OP50(xu363)* competent bacteria were transformed with dsRNA expression plasmids, which were extracted from the respective HT115 bacterial strains. The RNAi bacteria were grown in LB medium supplemented with 100 µg/mL ampicillin at 37 C° to reach log phase, washed with fresh medium, and concentrated 5-fold. Bacteria was then spread on RNAi plates, which are NGM plates containing 100 µg/mL ampicillin and 0.4 mM isopropyl ß-D-1-thiogalactopyranoside (IPTG). dsRNA-expressing bacteria were grown on plates at room temperature for two days. RNAi was initiated by letting the animals feed on the desired RNAi bacteria. Luciferase (L4440::luc, i.e., luci) RNAi vector was used as a non-targeting control in all experiments.

### Keio library screen

The *rnp-6(dh1127);Y41C4A.32(syb2725)* reporter strain was used for screening. The *E. coli* Keio knockout collection comprised two independent deletion strains for each of the 3985 nonessential genes, yielding a total of 7970 strains. All strains were tested as follows: briefly, Keio library bacteria were inoculated in 96-well plates overnight in LB (+ kanamycin) medium and seeded on 3 cm NGM plates. After two days of growth, ∼30 synchronized worm eggs (generated by bleaching) were seeded on the plates. The worms were scored after three days. For the primary screen, the plates were checked manually under a Leica stereo microscope (Leica M165 FC, LAS X). Mutant bacterial strains that resulted in more than 3 worms showing a bright mNeonGreen signal were marked and selected for further validation. In the confirmation screen, OP50 and the parental BW25113 strain were used as positive and negative control, respectively. Images of worms were captured for fluorescence intensity quantification. Bacterial hits were confirmed by Sanger sequencing.

### EMS mutagenesis screen

EMS mutagenesis was performed as described previously^22^. Briefly, ∼1,000 synchronized L4 larvae worms of *rnp-6(dh1127);Y41C4A.32(syb2725)* reporter strain were exposed to 0.15% ethyl methanesulfonate in M9 buffer for 4h at room temperature, and then washed and transferred to normal NGM plates for recovery. After overnight growth, P0 adult animals were transferred to new plates seeded with OP50 for egg laying. After 3 days of growing, adult F1 worms were bleached, and eggs were seeded onto NGM plates seeded with BW25113 bacteria. After 3- and 4-day growth, the plates were scored, and F2 worms that showed a bright fluorescence signal were singled to individual NGM plates. To exclude the false positive hits, the reporter fluorescence intensity of mutants growing on OP50 or BW25113 was measured, and the rescue efficiency (RE) of BW25113 diet was calculated. The mutants with RE value less than 0.5 were selected for further characterizations. The *rnp-6(dh1127);Y41C4A.32(syb2725)* animals were used as negative control in all the assays. To map causative mutations, Hawaiian-SNP mapping and whole genome sequencing were used as previously described^105^. In brief, EMS mutants were crossed with Hawaiian strain (CB4856) males. The non-fluorescent F1 worms were picked, raised until adulthood, and allowed to lay eggs on NGM plates seeded with BW25113. Fluorescent F2 adult worms were singled. After 5-days of growth, worms were then pooled together, and their genomic DNA purified using Gentra Puregene Kit (Qiagen). The pooled DNA was sequenced on an Illumina HiSeq platform (paired-end 150 nucleotide). MiModD pipeline (http://www.celegans.de/en/mimodd) was used to narrow down the causative mutations. The WBcel235/ce11 *C. elegans* assembly was used as a reference genome. Causative mutations were confirmed by multiple outcrosses.

### AID degradation experiments

The AID inducible-degradation system was designed following previously described protocols ^106^. Degron and GFP were inserted at the N-terminus of endogenous RNP-6. The transgenic line was crossed with TIR1-expression strain to generate the inducible-degradation strain AA5498 (*rnp-6*(*syb6010* [degron::3XFLAG::GFP::RNP-6]);Si57 [Peft-3::TIR1::mRuby::unc-54 3’UTR+ Cbr-unc-119(+)]). To induce RNP-6 degradation, strain AA5498 was grown on plates supplemented with different concentrations of Potassium Naphthaleneacetic Acid (K-NAA).

### RNA extraction and cDNA synthesis

*C. elegans* were lysed with QIAzol Lysis Reagent. RNA was extracted using chloroform extraction. Samples were then purified using RNeasy Mini Kit (Qiagen). Purity and concentration of the RNA samples were assessed using a NanoDrop 2000c (peqLab). cDNA synthesis was performed using iScript cDNA synthesis kit (Bio-Rad). Standard manufacturer protocols were followed for all mentioned commercial kits.

### RNA-Seq and bioinformatic analysis

1 µg of total RNA was used per sample for library preparation. The protocol of Illumina Tru-Seq stranded RiboZero was used for RNA preparation. After purification and validation (2200 TapeStation; Agilent Technologies), libraries were pooled for quantification using the KAPA Library Quantification kit (Peqlab) and the 7900HT Sequence Detection System (Applied Biosystems). The libraries were then sequenced with Illumina HiSeq4000 sequencing system using the paired-end 2×100 bp sequencing protocol. For data analysis, Wormbase genome (WBcel235_89) was used for alignment of the reads. Kallisto was used to map raw reads to reference transcriptome and quantify transcript abundance^107^. DESeq2 was used for each pairwise comparison^108^. For the splicing analysis, SAJR and IRfinder pipelines were used. The significant events from different pipelines were combined, and the unique events were kept for further analysis. Row Z-score heatmaps were generated by using the iHeatmap function from Flaski (Version 2.0.0) (DOI: 10.5281/zenodo.5254193). Adjusted p-value or q-value <0.05 is considered to be significant for gene expression and splicing. Gene enrichment visualization was performed with WormCat 2.0^24^.

### Alternative splicing PCR assay (RT-PCR)

A new batch of samples (different from the samples used for RNAseq) was used for RNA preparation and cDNA synthesis. Phusion Polymerase (Thermo Fisher) was used to amplify the *tos-1, tcer-1 and nhr-114* segments. PCR reactions were cycled 30 times with an annealing temperature of 53°C. RT-PCR products were visualized with ChemiDoc Imager (BioRad, ChemiDoc MP, Image Lab 6.1) after staining with Roti-GelStain (Carl Roth). Sequences of the primers used in the RT–PCR assays are found in Supplementary Table S15.

### Quantitative reverse transcription PCR (RT–qPCR)

Power SYBR Green master mix (Applied Biosystems) was used for RT-qPCR experiments. A JANUS automated workstation (PerkinElmer) was used for pipetting the reagents and cDNA samples into a 384 well plate. Thermal cycling was performed using a ViiA7 384 Real-Time PCR System machine (Applied Biosystems). *act-1* and *cdc-42* were used for internal normalization. Relative expression levels were calculated using the comparative CT method. Sequences of the primers used in the RT–qPCR assays are provided in Supplementary Table S15.

### Western blot

For *C. elegans* samples, animals were first washed with M9 buffer. Worm pellets were resuspended in 4% SDS buffer (4% SDS in 0.1 M Tris/HCl pH 8 with 1 mM EDTA) supplemented with cOmplete Protease Inhibitor (Roche) and PhosSTOP (Roche) and snap frozen in liquid nitrogen. Thawed samples were lysed using Bioruptor Sonication System (Diagenode), centrifuged (20,000g, 10 min) and protein concentrations were measured with Pierce BCA kit. Protein samples were then heated to 95 C° for 10 min in Laemmli buffer with 0.8% 2-mercaptoethanol in order to denature proteins. 10 ug protein samples were loaded on 4–15% Mini PROTEAN TGXTM Precast Protein Gels (Bio-Rad), and electrophoresis was performed at a constant voltage of 200V for around 40 min. After separation, the proteins were transferred to PVDF membranes using Trans-Blot TurboTM Transfer System (BioRad). 5% bovine serum albumin (BSA) or 5% milk in Tris-buffered Saline and Tween20 (TBST) were used for blocking of the membranes. After antibody incubations (anti-HA 1:1000, anti-Phospho-AMPKα (Thr172) 1:2000, anti-beta Actin 1:5000, Anti-Mouse HRP 1:5000, Anti-Rabbit HRP 1:5000 and Anti-Rat HRP 1:5000) and washing with TBST buffer, imaging of the membranes was performed with ChemiDoc Imager (BioRad, ChemiDoc MP, Image Lab 6.1). Western Lightning Plus Enhanced Chemiluminescence Substrate (PerkinElmer) was used as the chemiluminescence reagent. A list of antibodies is provided in Supplementary Table S15.

### Metabolomics and lipidomics sample preparation

Worms were synchronized by egg laying and collected when they reached the young adult stage. For each sample, ∼2000 worms were washed three times with ddH₂O, snap-frozen in liquid nitrogen, and stored at −80°C before use. For the metabolite/lipid extraction, 500 µL extraction buffer (MTBE:MeOH:H2O, 50:30:20) supplemented with internal standards was added to each sample. Worm pellets were homogenized with ∼100 µL 1 mm zirconia beads using a TissueLyser (Qiagen) at 50 Hz and 4°C for 20 min. After initial homogenization, 500 µL extraction buffer was added to each sample and homogenization resumed for 5 min. Worm lysates were centrifuged at 21000 x g and 4°C for 10 min. Supernatant was transferred to new tubes. Residual buffer was removed before drying protein pellets under a fume hood overnight. Protein concentration was determined using a BCA kit (ThermoFisher). The cleared supernatant was mixed with 200 µL MTBE (Sigma) and 150 µL H2O, and incubated at 15°C for 10 min. Samples were centrifuged at 15°C and 16000 x g for 10 min to obtain phase separation (top lipid phase, bottom polar metabolites phase). 650 µL of the lipid phase was transferred to new tubes and dried in a SpeedVac concentrator at 20°C and 1000 rpm for ∼2 h. 600 µL polar phase solution was transferred to new tubes and dried in a Speed Vac concentrator at 20°C and 1000 rpm for ∼6 h. Samples were stored at –80°C until further processing.

### Semi-targeted liquid chromatography-high-resolution mass spectrometry-based (LC-HRS-MS) analysis of amine-containing metabolites

Amine-containing compounds were analyzed using a Q-Exactive Plus high-resolution mass spectrometer coupled to a Vanquish UHPLC chromatography system (Thermo Fisher Scientific). As previously described, dried extracts were reconstituted in 150 µL LC-MS-grade water at 4°C and 1500 rpm shaking for 10 min. After centrifugation, 50 µL supernatant was mixed with 25 µL 100 mM sodium carbonate, followed by 25 µL 2% (v/v) benzoyl chloride in acetonitrile (UPLC/MS-grade, Biosolve) ^109^. Samples were mixed and stored at 20°C until analysis. 1 µL of derivatized sample was injected onto a 100 × 2.1 mm HSS T3 UPLC column (Waters) at 40°C, 400 µL/min, using a binary buffer system: buffer A (10 mM ammonium formate, 0.15% formic acid in water) and buffer B (acetonitrile). LC gradient: 0% B (0 min); 0–15% B (0–4.1 min); 15–17% B (4.1–4.5 min); 17–55% B (4.5–11 min); 55–70% B (11–11.5 min); 70–100% B (11.5–13 min); 100% B (13–14 min); 100–0% B (14–14.1 min); 0% B (14.1–19 min). MS acquisition was performed in positive ionization mode (m/z 100–1000), with the following source settings: 3.5 kV spray voltage, capillary temperature 300°C, sheath gas 60 AU, aux gas 20 AU at 330°C, sweep gas 2 AU, RF lens 60. Raw mass spectra files were converted to mzML using MSConvert (v3.0.22060) ^110^, and analyzed in El Maven (v0.12.0) ^111^. Area of the protonated [M + nBz + H]^+^ mass peaks of every required compound was extracted and integrated (mass accuracy of <5 ppm, retention time tolerance of <0.05 min compared to reference compounds). Data were normalized to internal standards and protein content.

### Anion-exchange chromatography mass spectrometry (AEX-MS) for the analysis of anionic metabolites

Extracted metabolites were reconstituted in 150 µL UPLC/MS grade water (Biosolve) of which 100 µL was transferred to polypropylene autosampler vials (Chromatography Accessories Trott). Analysis was performed as previously described, using a Dionex Integrion ion chromatography system (Thermo Fisher Scientific)^4^. 5 µL of extract was injected (push-full mode, overfill factor 1) onto a Dionex IonPac AS11-HC column (2 × 250 mm, 4 µm particle size) equipped with a AG11-HC guard column (2 × 50 mm, 4 µm) at 30°C and autosampler at 6°C. Metabolites were separated at 380 µL/min using a KOH cartridge (Eluent Generator, Thermo Scientific) with the following gradient: 0–3 min, 10 mM; 3–12 min, 10–50 mM; 12–19 min, 50–100 mM; 19–22 min, 100 mM; 22–23 min, 100–10 mM; re-equilibration at 10 mM (3 min). Eluting metabolites were detected in negative ion mode (m/z 77–770) on a Q-Exactive HF MS. Source settings: 3.2 kV spray voltage, capillary 300°C, sheath gas 50 AU, aux gas 14 AU at 380°C, sweep gas 3 AU, S-lens 40. Raw mass spectra files were converted to mzML using MSConvert (v3.0.22060)^3^, and analyzed in El Maven (v0.12.0)^2^. Area of the deprotonated [M-H^+^]^-1^ or doubly deprotonated [M-2H]^-2^ isotopologues mass peaks of every required compound was extracted and integrated (mass accuracy of <5 ppm, retention time tolerance of <0.05 min compared to reference compounds). Data were normalized to internal standards and protein content.

### Liquid Chromatography-High Resolution Mass Spectrometry-based (LC-HRMS) analysis of lipids

Stored (-80°C) lipid extracts were reconstituted in 150 µL UPLC-grade acetonitrile:isopropanol (70:30, v/v). After vortexing, samples were incubated for 10 min at 4°C with continuous shaking. Samples were clarified by centrifugation at 16000 x g, 4°C, for 5 min. Supernatants were transferred to 2 mL glass vials equipped with 300 µL glass inserts (Chromatography Zubehör Trott). Aliquots of 20 µL from each sample were pooled to generate quality control (QC) samples, which were injected after every 10th sample or after each replicate group. Samples and QCs were maintained at 6°C in a Vanquish UHPLC system (Thermo Fisher Scientific) fitted with a quaternary pump and coupled to a TimsTOF Pro 2 HRMS with a heated ESI (VIP-HESI) source (Bruker Daltonics). For each run, 1 µL sample was injected onto a 100 x 2.1 mm CSH C18 UPLC column (1.7 µm, Waters). The chromatographic gradient was performed at 400 µL/min using buffer A (10 mM ammonium formate, 0.1% formic acid in acetonitrile:water, 60:40, v/v) and buffer B (10 mM ammonium formate, 0.1% formic acid in isopropanol:acetonitrile, 90:10, v/v) as follows: 0–0.5 min, 45–48% B; 0.5–1 min, 48–55% B; 1–1.8 min, 55–60% B; 1.8–10 min, 60–85% B; 10–11 min, 85–99% B; 11–11.5 min, 99% B; 11.7–15 min, re-equilibration at 45% B (total run time: 15 min/sample). Prior to each batch, mass and mobility calibrations were performed using a 1:1 mixture of 10 mM sodium formate and ESI-L Low Concentration Tuning Mix (Agilent). Data were acquired in data-dependent PASEF mode, primarily in positive ionization mode (source settings: 4.5 kV capillary, 500 V end plate offset, nebulizer 2 bar, dry gas 8 L/min at 230°C, sheath gas 4 L/min at 400°C). For MS/MS, isolation width was set to 2 mD and collision energy to 30 eV. Pooled QC samples were additionally injected in negative mode (collision energy 40 eV, - 3.5 kV capillary) for further fatty acid annotation. Samples were analyzed in randomized order. QC samples were analyzed after every 10th injection in both positive and negative ionization modes to ensure data quality and stability. Raw spectra were processed in MetaboScape (v2024) to extract features and annotate lipid species, using pooled QC samples for validation. Lipids were only included in downstream analysis if their abundance in QC samples was at least threefold higher than in extraction blanks.

### Human sample handling and preparation

The peripheral blood samples of VRJS patients were collected in the Center for Rare Diseases, University Hospital of Cologne, Germany. All patients and/or custodians gave informed consent according to local institutional review board approval (20-1711, Medical Faculty at University of Cologne). The plasma was isolated with standard protocol and processed for metabolomic and lipidomic analysis in the Metabolomic Core Facility of the Max Planck Institute for Biology of Ageing.

### Statistics & Reproducibility

In all figures, the numbers of independent replicates and the total number of animals analyzed are indicated in each panel. All statistical analyses were performed in GraphPad Prism (Version 9.0.0 (86)). Asterisks denote corresponding statistical significance *P < 0.05; **P < 0.01; ***P < 0.001; ****P < 0.0001. Data distribution was assumed to be normal but this was not formally tested. No statistical method was used to predetermine sample size but our sample sizes are similar to those reported in previous publications^23,73,112,113^. Missing or outlier data points were excluded from the analyses. At least three independent experiments for each assay were performed to verify the reproducibility of the findings, unless indicated otherwise. For worm experiments, samples preparation and data collection were randomized. For lifespan experiments, all the genotypes were blinded before the assays. For cold tolerance, developmental rate, body size, brood size, Western blot, and imaging experiments, the genotypes were not blinded before assay, as mutant worms have obvious phenotypes that revealed the sample identity (body size and developmental rate). However, worms were randomly picked and assigned to the different treatment conditions in a random order. For RNA-seq experiments, the genotypes were not blinded before collecting samples. Once the RNA samples were ready, they were processed at the Cologne Center for Genomics (CCG) in a blinded manner.

## Data availability

The RNA-seq datasets generated in this study are available in the GEO datasets with the accession number GSE307065 as of the date of publication. The list of *nhr-114* target genes was as defined^61^, available under the accession number GSE211747. The siPUF60-treated RNAseq data was retrieved under accession number OEP004324^78^ in NODE (The National Omics Data Encyclopedia) database.

## Acknowledgements

We thank the Caenorhabditis Genetics Center (University of Minnesota), Lianfeng Wu (Westlake University), and William Mair (Harvard T.H. Chan School of Public Health) for kindly providing *C. elegans* strains. We thank the Bioinformatics, Imaging, Metabolomics, and Proteomics core facilities of the Max Planck Institute for Biology of Ageing for their technical support. We would also like to thank the members of the Antebi lab, especially Dr. Kreuz, Dr. Tabrez and Dr. Kawamura for valuable comments on the manuscript. We also thank Dr. Filipe Cabreiro for his valuable feedback on the manuscript. We thank the *PostDoc* Seed funding 2023 (W.H.) from CECAD Cluster for Excellence, University of Cologne (Gefördert durch die Deutsche Forschungsgemeinschaft (DFG) im Rahmen der Exzellenzstrategie des Bundes und der Länder - EXC 2030 – 390661388), Cologne Graduate School for Ageing Research (J.K.) and the Max Planck Society, Germany (A.A.) for funding this project.

## Author contributions

W.H., J.K. and A.A. conceived and designed the study. W.H., J.K. and A.L. performed the investigation in *C. elegans*. K.C. performed the western blot experiment. W.H. and J.K. analyzed the data. D.H. and E.B. collected, processed, and analyzed patient samples. W.H. wrote the original draft of the manuscript; W.H., J.K. and A.A. reviewed and edited the final version. A.A. acquired funding for the project.

