## Supplementary_figures for "Vitamin B_12_ alleviates Verheij syndrome-like defects via phospholipid remodeling in a *C. elegans* PUF60 spliceosomopathy model"

Figure S1

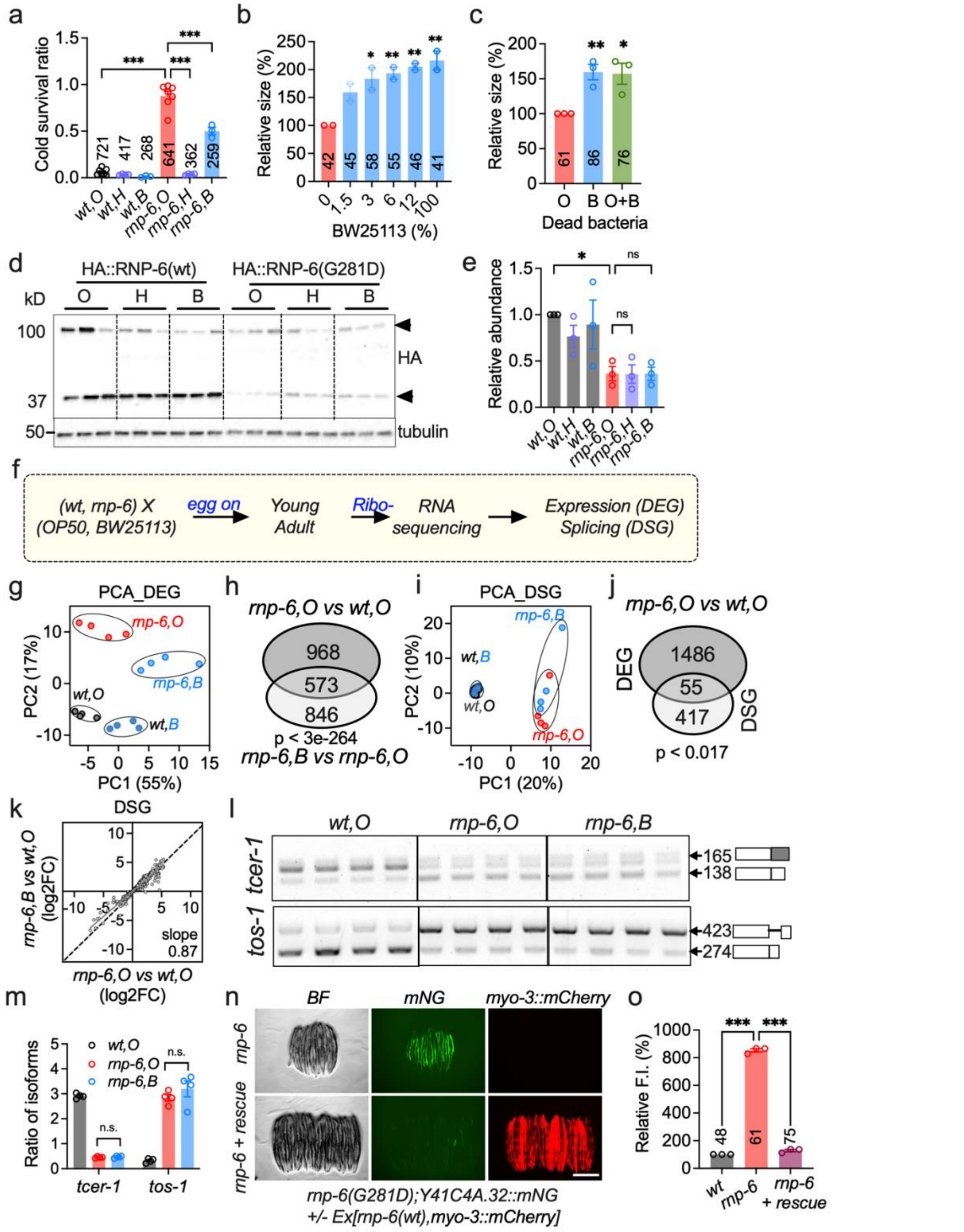


**Figure S1. A bacterial *E. coli* K12 diet mitigates developmental defects in an *rnp-6*/VRJS *C. elegans* model. a**, Assessment of cold tolerance in worms fed *E. coli* K12 strains; bars represent the ratio of alive to dead worms per genotype and diet after cold shock ($n$ = 3–7). **b**, Evaluation of BW25113 dilution effects on worm body size ($n$ = 2). **c**, Body size of *rnp-6* mutant worms fed UV-killed BW25113 and OP50-BW25113 (ratio 1:1) mixture ($n$ = 3). **d–e**, Total RNP-6 protein abundance in response to K12 diets. The total amounts of two predominant RNP-6 isoforms (as denoted by arrowheads) were quantified (*n* = 3). **f**, Overview of RNA sequencing experimental design. Ribo-, ribosomal RNA depletion. **g**, Principal component analysis of differentially expressed genes (DEG), four biological replicates were analyzed for each condition. **h**, Venn diagram of DEGs changed by *rnp-6* mutation on OP50 and BW25113. **i**, Principal component analysis of differentially spliced genes (DSG). **j**, Overlap between DEGs and DSGs, shown by Venn diagram. **k**, Effect of BW25113 *E. coli* on alternative splicing; shown are 638 events significantly altered by *rnp-6(G281D)* on OP50. l–**m**, RT-PCR splicing analyses of *tos-1* and *tcer-1* (*n* = 4). **n**–**o**, Effect of *rnp-6(wt)* overexpression on Y41C4A.32::mNG reporter expression. *myo-3::mCherry* co-injection marker indicates transgenic animals (*n* = 3, scale bar: 500 µm). Data are presented as mean ± s.e.m. unless otherwise indicated; "*n*" denotes experimental replicates, with animal counts summarized in each respective bar. Statistical significance was determined using one-way ANOVA with Dunnett's multiple comparisons (**a**, **b**, **c**, **e**, **m**, **o**). All the source data for figures are provided in supplementary tables.

Figure S2

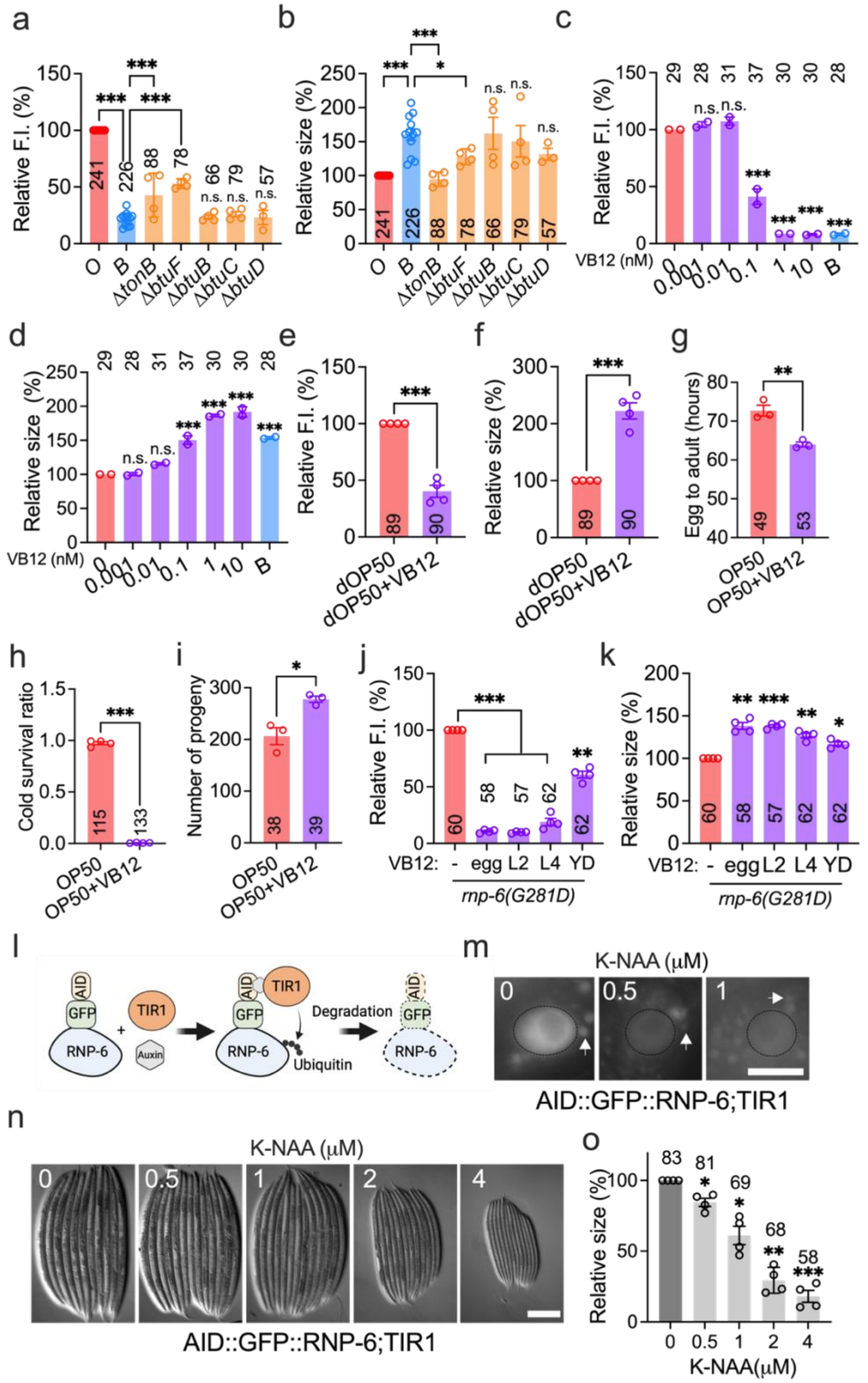


**Figure S2. A complementary two-way genetic screen identifies vitamin B12 as an alleviator of *rnp-6(G281D)* mutant defects. a–b**, Effects of alternative vitamin B12 transporters on Y41C4A.32 reporter expression (**a**) and body size (**b**) in *rnp-6(G281D)* mutants ($n$ = 4–12). **c–d**, Dose-response effects of vitamin VB12 supplementation on *rnp-6(G281D)* phenotypes ($n$ = 2). **e–f**, Impact of VB12 supplementation in *rnp-6(G281D)* mutants fed UV-killed OP50 ($n$ = 4). **g–i**, Effects of VB12 supplementation on developmental rate (**g**, *n* = 3), cold tolerance (**h**, *n* = 4), and brood size (**i**, *n* = 3). **j–k**, Effects of VB12 supplementation administered from distinct developmental stages on Y41C4A.32 reporter expression (**j**) and body size (**k**) in *rnp-6(G281D)* mutants (*n* = 4). Worms were imaged on day 2 of adulthood. **l**, Schematic illustrating the RNP-6 inducible degradation system. The image was made in BioRender. **m**, Effects of K-NAA supplementation on GFP::RNP-6 fluorescence intensity on OP50 (scale bar: 10 μm). The nuclei of the intestine are indicated with circles. The arrows indicate gut autofluorescent granules. **n–o**, Impact of various K-NAA concentrations on worm body size on OP50 (*n* = 4, scale bar: 200 μm). The final concentration of VB12 is 1 nM unless otherwise noted. Statistical significance was determined using one-way ANOVA with Dunnett's multiple comparisons. Significance is represented as *P < 0.05, **P < 0.01, ***P < 0.001, n.s.: not significant.

Figure S3

a

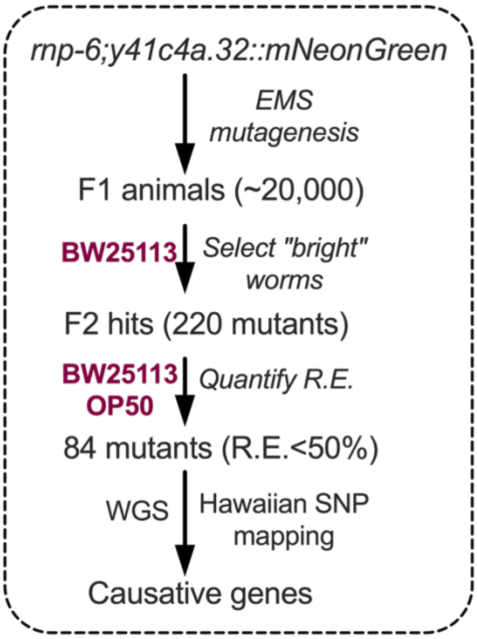


**Figure S3. Vitamin $B_{12}$ alleviates *rnp-6(G281D)* mutant defects via methionine metabolism pathways. a**, Schematic of EMS mutagenesis screen for host factors. R.E. (rescue efficacy) = (Fluorescent intensity on OP50 - Fluorescent intensity on BW25113)/ Fluorescent intensity on OP50.

Figure S4

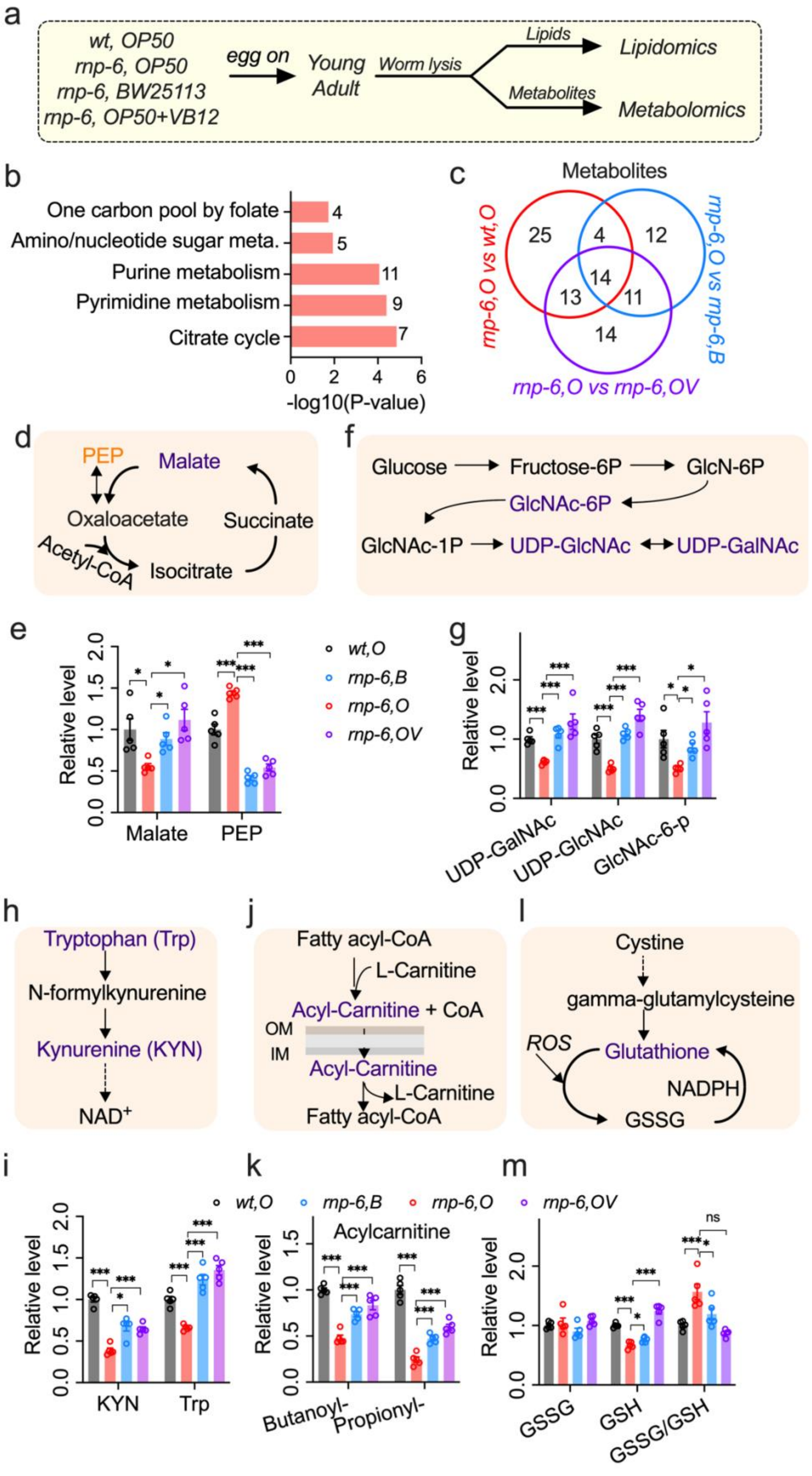
a
wt, OP50
rnp-6, OP50
rnp-6, BW25113
rnp-6, OP50+VB12
egg on
Young Adult
Worm lysis
Lipids
Lipidomics
Metabolites
Metabolomics
b
One carbon pool by folate
Amino/nucleotide sugar meta.
Purine metabolism
Pyrimidine metabolism
Citrate cycle
-log10(P-value)
c
Metabolites
rnp-6,O vs wt,O
rnp-6,O vs rnp-6,B
rnp-6,O vs rnp-6,OV
d
PEP
Malate
Oxaloacetate
Succinate
Acetyl-CoA
Isocitrate
e
Relative level
Malate
PEP
wt,O
rnp-6,B
rnp-6,O
rnp-6,OV
f
Glucose
Fructose-6P
GlcN-6P
GlcNAc-6P
GlcNAc-1P
UDP-GlcNAc
UDP-GalNAc
g
Relative level
UDP-GalNAc
UDP-GlcNAc
GlcNAc-6-p
h
Tryptophan (Trp)
N-formylkynurenine
Kynurenine (KYN)
NAD+
j
Fatty acyl-CoA
L-Carnitine
Acyl-Carnitine + CoA
OM
IM
Acyl-Carnitine
L-Carnitine
Fatty acyl-CoA
l
Cystine
gamma-glutamylcysteine
ROS
Glutathione
NADPH
GSSG
i
Relative level
KYN
Trp
k
Acylcarnitine
Relative level
Butanoyl-
Propionyl-
m
Relative level
GSSG
GSH
GSSG/GSH
ns
wt,O
rnp-6,B
rnp-6,O
rnp-6,OV

**Figure S4. Vitamin $B_{12}$ restores methylation potential and phosphatidylcholine metabolism in *rnp-6* mutants. a**, Workflow for metabolomic and lipidomic analyses. Five replicates per condition. **b**, MetaboAnalyst-based pathway analysis of metabolites significantly altered in *rnp-6(G281D)* mutants. **c**, Venn diagram of significantly altered metabolites. Simplified schematics and abundance changes are shown for the following pathways: **d–e**, tricarboxylic acid (TCA) cycle; **f–g**, amino sugar and nucleotide sugar metabolism; **h–i**, kynurenine pathway; **j–k**, mitochondrial fatty acid metabolism; **l–m**, glutathione metabolism. Statistical significance was determined using one-way ANOVA with Dunnett's multiple comparisons. Significance is represented as $^{*}P < 0.05$, $^{**}P < 0.01$, $^{***}P < 0.001$, n.s.: not significant.

Figure S5

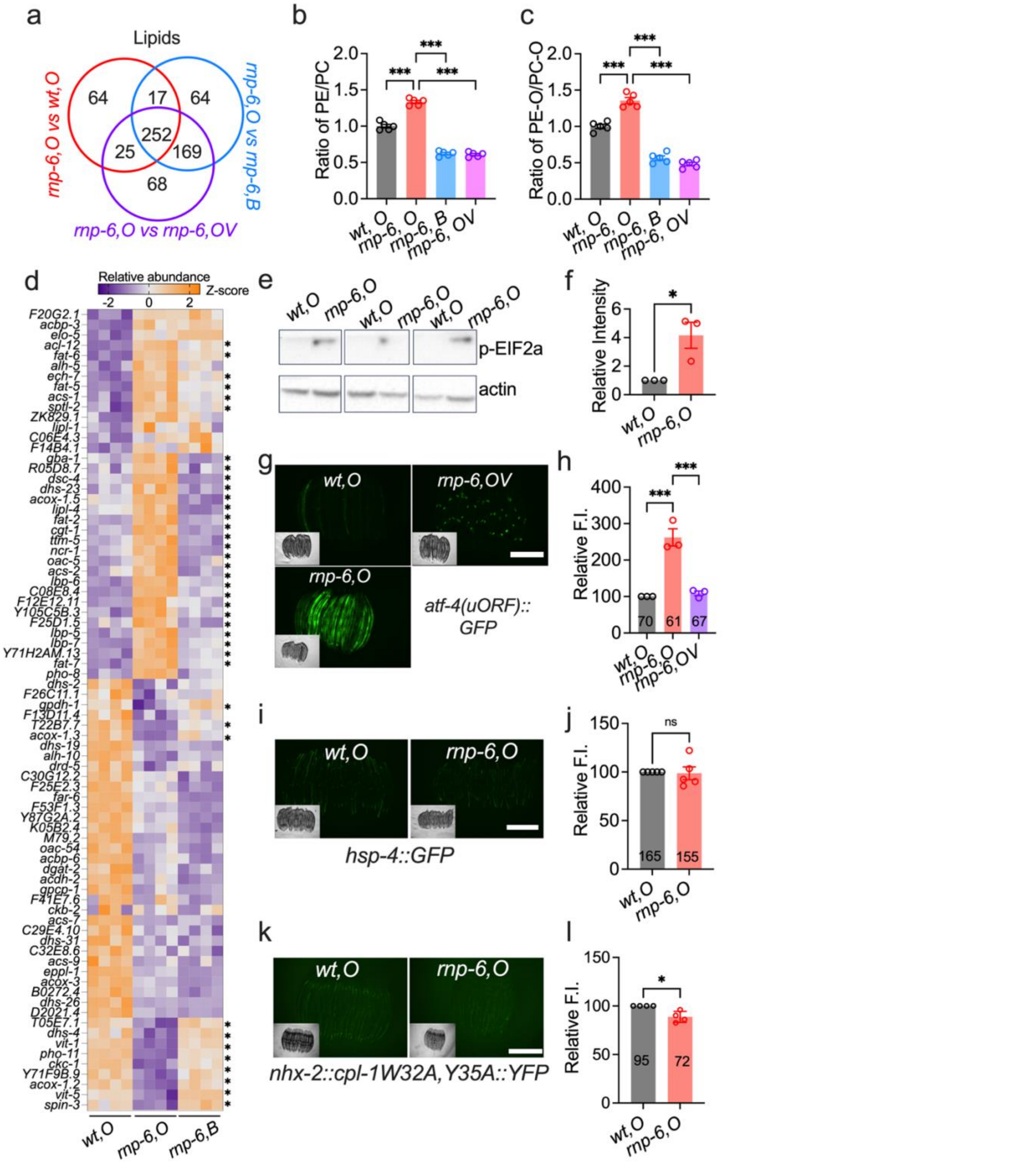


**Figure S5. Vitamin B$_{12}$ restores phosphatidylcholine metabolism and the integrated stress response in *rnp-6* mutants. a**, Venn diagram of significantly altered lipid species. **b**, Ratio of phosphatidylethanolamine (PE) to phosphatidylcholine (PC) changes. **c**, Ratio of ether-linked PE (PE-O) to ether-linked PC (PC-O) changes. **d**, Heat map of differentially expressed genes (DEGs) associated with lipid metabolism; "*" denotes genes significantly restored by BW25113 diet. **e–f**, Western blot analysis of eIF2α phosphorylation in *rnp-6*

mutants, with three biological replicates analyzed (n = 3). **g–h**, Representative images and quantification of ISR reporter strain atf-4(uORF)::GFP expression (scale bar: 500 μm). **i–j**, Representative images and quantification of unfolded protein response of the endoplasmic reticulum ($UPR^{ER}$) reporter strain hsp-4::GFP expression (n = 5, scale bar: 500 μm). **k–l**, Representative images and quantification of CPL-1 reporter strain cpl-1W32A,Y35A::YFP expression (*n* = 4, scale bar: 500 μm). Statistical significance was determined using unpaired t-test (**f**, **j**, **l**) and one-way ANOVA with Dunnett's multiple comparisons (**h**). Significance is represented as *P < 0.05, **P < 0.01, ***P < 0.001, n.s.: not significant.

Figure S6

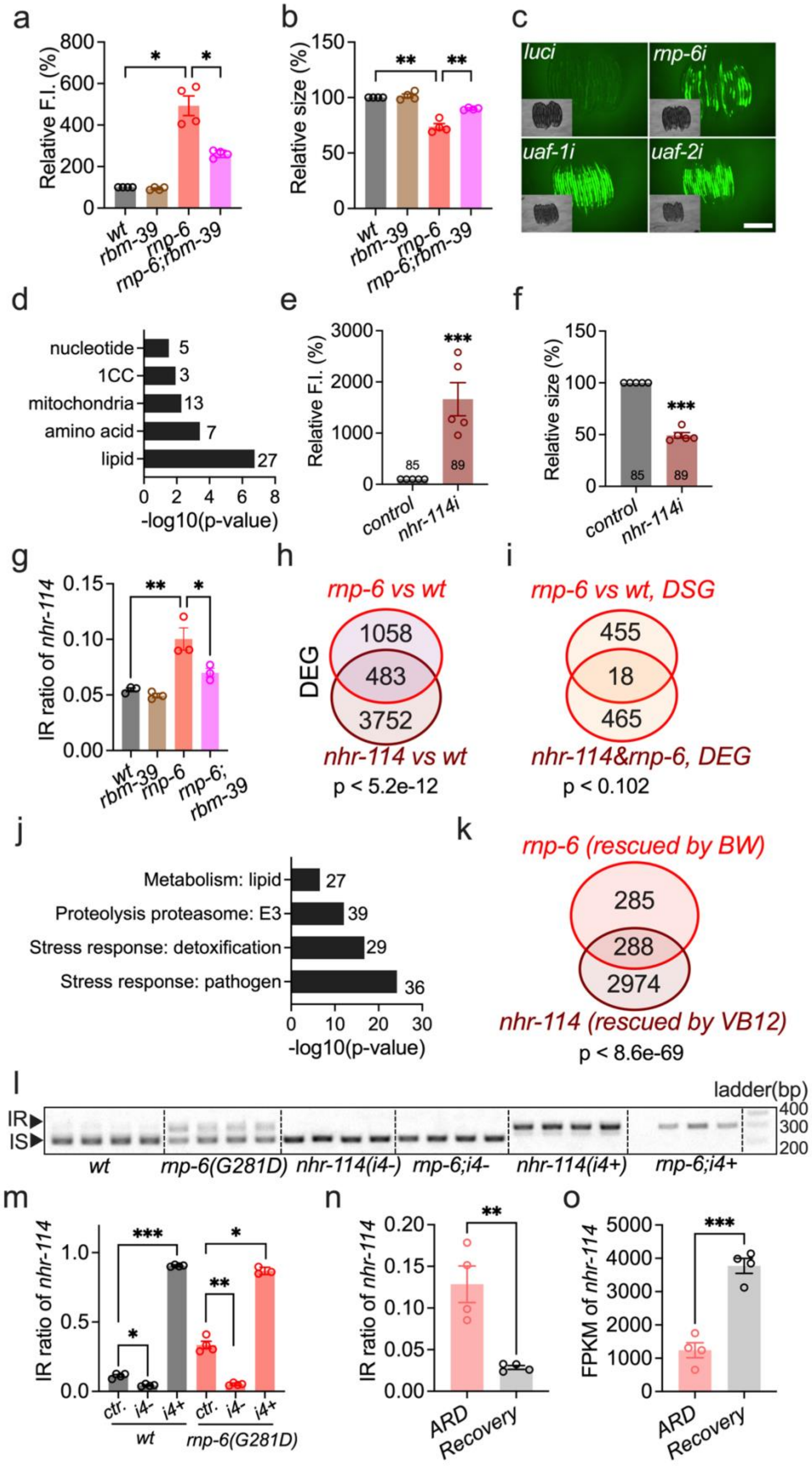

a
Relative F.I. (%)
wt
rbm-39
rnp-6
rnp-6;rbm-39
b
Relative size (%)
c
luci
rnp-6i
uaf-1i
uaf-2i
d
nucleotide
1CC
mitochondria
amino acid
lipid
-log10(p-value)
e
Relative F.I. (%)
control
nhr-114i
f
Relative size (%)
g
IR ratio of nhr-114
h
rnp-6 vs wt
DEG
nhr-114 vs wt
p < 5.2e-12
i
rnp-6 vs wt, DSG
nhr-114&rnp-6, DEG
p < 0.102
j
Metabolism: lipid
Proteolysis proteasome: E3
Stress response: detoxification
Stress response: pathogen
-log10(p-value)
k
rnp-6 (rescued by BW)
nhr-114 (rescued by VB12)
p < 8.6e-69
l
ladder(bp)
IR
IS
wt
rnp-6(G281D)
nhr-114(i4-)
rnp-6;i4-
nhr-114(i4+)
rnp-6;i4+
m
IR ratio of nhr-114
ctr.
i4-
i4+
n
IR ratio of nhr-114
ARD
Recovery
o
FPKM of nhr-114

**Figure S6. *rnp-6* mutant defects are primarily caused by aberrant splicing of transcription factor *nhr-114*. a–b**, Quantification of Y41C4A.32::mNG fluorescent intensity (**a**) and worm body size (**b**) in the *rbm-39(S294L)* mutant background ($n = 4$). **c**, Representative images of Y41C4A.32::mNG fluorescence intensity and body size following *uaf-1* and *uaf-2* RNAi treatment. The *mthf-1(W601stop);Y41C4A.32::mNG* strain was used to mitigate suppression from HT115 on Y41C4A.32::mNG reporter expression, with luciferase-knockdown used as a control (scale bar: 500 μm). All images were adjusted for a better view with identical settings. **d**, Gene set enrichment analysis of differentially spliced genes involved in metabolism, performed with WormCat 2.0. **e–f**, Quantification of Y41C4A.32::mNG fluorescence intensity (**e**) and worm body size (**f**) after *nhr-114* RNAi (n = 5) in wild-type N2 background. OP50 RNAi bacteria was used to mitigate the suppression effects from HT115. **g**, RNAseq analysis of *nhr-114* intron 4 retention. **h**, Venn diagram illustrating differentially expressed genes in the *nhr-114* deletion mutant and the *rnp-6* mutant. **i**, Venn diagram illustrating genes that are differentially spliced. **j**, Gene set enrichment analysis of the overlapped genes in (h). **k**, Venn diagram of differentially expressed genes significantly rescued by either VB12 supplementation in *nhr-114* deletion mutants or BW25113 diet in *rnp-6* mutants. **l–m**, RT-PCR analysis of *nhr-114* intron 4 retention. *nhr-114(+)* results in almost complete intron retention; one biological replicate for *rnp-6; nhr-114(i4+)* was excluded from quantification due to technical failure (n = 3–4). **n–o**, Quantification of *nhr-114* intron 4 retention and expression levels in ARD worms, analyzed by RNA-seq (n = 4). Statistical significance was determined using unpaired t-test (**e**, **f**, **n**, **o**) and one-way ANOVA with Dunnett's multiple comparisons (**a**, **b**, **g**, **m**). Significance is represented as $*P < 0.05$, $**P < 0.01$, $***P < 0.001$, n.s.: not significant.

Figure S7

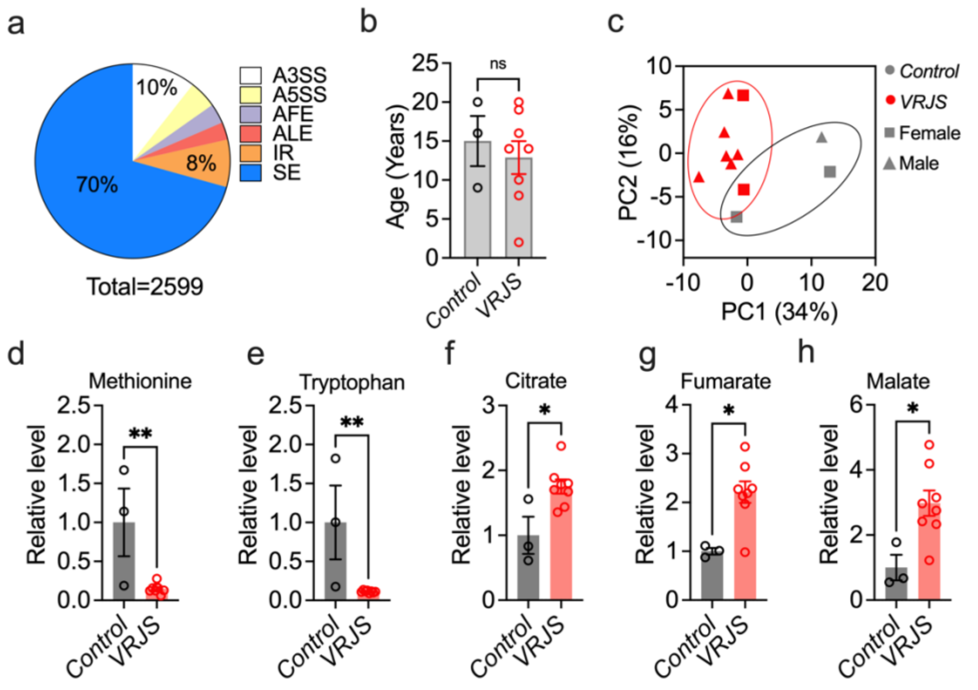


**Figure S7. PUF60 deficiency impairs 1CC and phospholipid metabolism in human cells.** **a,** Pie chart displaying differentially spliced events in the siPUF60-treated PC9 cell line. **b**, Age distribution of individuals for blood samples. **c**, PCA analysis of metabolomic and lipidomic data. **d-h**, The relative abundance of methionine (d), tryptophan (e), citrate (f), fumarate (g) and malate (h) in VRJS patient plasma. Statistical significance was determined using unpaired t-test  Significance is represented as *P < 0.05, **P < 0.01, ***P < 0.001, n.s.: not significant.

Figure S8

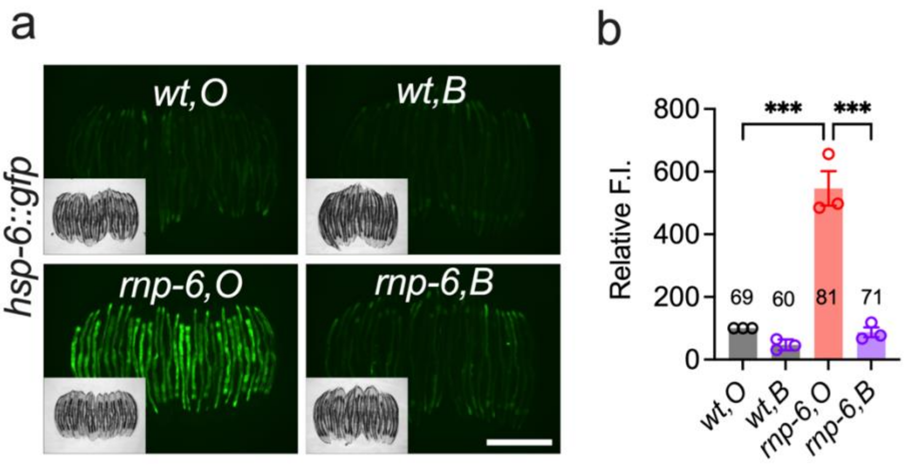


**Figure S8. *E. coli* BW25113 alleviates mitochondrial stress in *rnp-6(G281D)* mutants. a,** Representative images of *hsp-6::gfp* expression in *rnp-6* mutants. Scale bar, 500 µm. n=3. b, Quantification of *hsp-6::gfp* fluorescent intensity. One-way ANOVA with Dunnett's multiple comparison test was used for statistical analysis. ***P < 0.001.
